# DNA-sequence and epigenomic determinants of local rates of transcription elongation

**DOI:** 10.1101/2023.12.21.572932

**Authors:** Lingjie Liu, Yixin Zhao, Adam Siepel

## Abstract

Across all branches of life, transcription elongation is a crucial, regulated phase in gene expression. Many recent studies in eukaryotes have focused on the regulation of promoter-proximal pausing of RNA Polymerase II (Pol II), but rates of productive elongation also vary substantially throughout the gene body, both within and across genes. Here, we introduce a probabilistic model for systematically evaluating potential determinants of the local elongation rate based on nascent RNA sequencing (NRS) data. Our model is derived from a unified model for both the kinetics of Pol II movement along the DNA template and the generation of NRS read counts at steady state. It allows for a continuously variable elongation rate along the gene body, with the rate at each nucleotide defined by a generalized linear relationship with nearby genomic and epigenomic features. High-dimensional feature vectors are accommodated through a sparse-regression extension. We show with simulations that the model allows accurate detection of associated features and accurate prediction of local elongation rates. In an analysis of public PRO-seq and epigenomic data, we identify several features that are strongly associated with reductions in the local elongation rate, including DNA methylation, splice sites, RNA stem-loops, CTCF binding sites, and several histone marks, including H3K36me3 and H4K20me1. By contrast, low-complexity sequences and H3K79me2 marks are associated with increases in elongation rate. In an analysis of DNA *k*-mers, we find that cytosine nucleotides are strongly associated with reductions in local elongation rate, particularly when preceded by guanines and followed by adenines or thymines. Increases in elongation rate are associated with thymines and A+T-rich *k*-mers. These associations are generally shared across cell types, and by considering them our model is effective at predicting features of held-out PRO-seq data. Overall, our analysis is the first to permit genome-wide predictions of relative nucleotide-specific elongation rates based on complex sets of genomic and epigenomic covariates. We have made predictions available for the K562, CD14+, MCF-7, and HeLa-S3 cell types in a UCSC Genome Browser track.

## Introduction

An enduring challenge in the study of eukaryotic gene regulation is that there is no single, well-defined point of control for gene expression. Instead, rates and patterns of expression are influenced at a broad array of cellular stages, ranging from pre-transcriptional chromatin remodeling to transcriptional, post-transcriptional, translational and post-translational steps. Even within the critical stage of transcription—where research has traditionally focused on control of transcription initiation—many different steps can be regulated.

After RNA Polymerase II (Pol II) has been recruited to a promoter, and together with its cofactors, unwound the DNA and established a stable RNA-DNA hybrid, it begins to translocate along the DNA template and synthesize a nascent RNA molecule [1, 2]. This process of “productive elongation” is not perfectly homogenous but occurs at variable rates along the DNA template, sometimes pausing entirely for minutes at a time. A particular focus of recent research has been the tendency of Pol II to exhibit a pronounced pause ∼20–60 bp downstream of the TSS. It has become evident that such promoter-proximal pausing is remarkably widespread, both across metazoan species and across genes, and escape from such pausing appears to be regulated in many cases [3, 4]. A great deal of attention has been devoted to working out the molecular mechanisms underlying this process.

Rates of elongation also vary throughout the gene body, but the determinants of this less stereotypical rate variation are less well understood. What is known is the following. First, productive elongation rates vary considerably across genes. In mammals, elongation through gene bodies occurs at an average rate of roughly 2 kb/min but this rate can vary by fourfold or more across genes, and it can also vary considerably for the same gene across cell types or conditions [5–10]. Second, the local elongation rate changes along each individual gene body, tending to increase with distance from the TSS but becoming reduced, again, at exons and near the termination site [5, 6]. Third, average elongation rates for genes are correlated with a wide variety of genomic and epigenomic features, including G+C content, exon density, nucleosome density, DNA methylation, histone marks such as H3K4me1, H3K4me3, H3K36me3, H3K79me2, and H4K20me1, stability of the DNA-RNA hybrid, the density of low-complexity sequences, as well as various DNA 5-mer frequencies [5–7, 9, 11, 12] (reviewed in [4, 13–15]). Fourth, elongation rates are also positively correlated with the Pol II density itself, suggesting that the activity of one polymerase somehow facilitates the progress of others (perhaps through phenomena such as gradual loss of pausing factors, chromatin remodeling, or impact on DNA torsion) [5]. Elongation rates are similarly positively correlated with gene length [7]. Finally, in at least some cases, it is clear that elongation rates are not only indirectly influenced by structural features of DNA, RNA, or chromatin, but are actively regulated in response to various cellular stimuli, with dysregulation of elongation rates potentially contributing to disease progression [14].

Other studies have focused specifically on pausing of Pol II. Aside from the pronounced pausing that occurs proximal to the promoter, many, typically more subtle, pause sites occur within gene bodies, and, in the aggregate, these sites have a major effect on the dynamics of transcription elongation [16–19] (see also [20] for a recent study in yeast). These gene-body pause sites replicate well across experiments but vary substantially in their density across genes; they also occur in divergent antisense transcripts and enhancer RNAs as well as in gene bodies [18]. Such pausing has been reported to be associated with intron-exon boundaries, alternative splicing, certain properties of DNA shape and the RNA-DNA hybrid, DNA methylation, binding sites for factors such as ESR1, PAX5, SMAD3, YY1, and CTCF, and particular sequence motifs, some of which are distinct from those associated with promoter-proximal pausing [18, 19, 21–23].

Despite these findings, much remains unclear about the determinants of local elongation rates through gene bodies. Most studies have either been based on the measurement of rates of progress of Pol II “waves” in time-course experiments [5–10], or on pre-identified gene-body pause sites [18, 19]. The wave experiments have limited genomic resolution, because waves tend to move tens of kilobases between time points, and are therefore better suited for evaluating correlates of average genic elongation rates than of local rates. They also are restricted to subsets of genes at which transcription can be induced or repressed, and to longer genes. The experiments based on pre-identified pause sites have better genomic resolution but tend to reflect only the most extreme reductions in rate—ones sufficient to produce a statistically significant local peak in nascent RNA sequencing read counts. Furthermore, with both types of studies, it is difficult to make sense of the observed genomic and epigenomic covariates because many of them are also strongly correlated with one another.

In this study, we revisit these questions using a fundamentally different statistical modeling approach. Our method is based on a recently developed “unified model” for nascent RNA sequencing (NRS) data, which describes both the kinetics of Pol II movement on the DNA template and the generation of NRS read counts [24, 25]. We adapt this model to allow for a continuously variable elongation rate along the genome, using a generalized linear model to capture the relationship between the local rate and nearby genomic features. In this way, we avoid a dependency on predefined pause-sites, and jointly consider the influences of a large and diverse collection of features on elongation rate. By accounting for differences across genes in initiation rate, we are able to efficiently pool information across all genes in the genome, and extract high-resolution information about relative local elongation rate from steady-state data. This strategy avoids a dependency on specially designed time-course experiments, and enables application to a wide variety of existing public data sets. We consider both epigenomic and DNA-sequence covariates of elongation rate, and after showing that our methods are effective with simulated data, we apply them to public data for four mammalian cell lines, identifying a number of both previously known and novel correlates of elongation rate. We then use our models to predict relative nucleotide-specific elongation rates genome-wide and make our predictions available in a UCSC Genome Browser track (available at https://bit.ly/elongation-rate-tracks).

## Results

### A generalized linear model for variable elongation rates

Our previous “unified model” describes both the stochastic movement of Pol II along a DNA template and the probabilistic generation of nascent RNA sequencing (NRS) read counts from underlying Pol II densities [24]. The model has two layers: a continuous-time Markov model for the movement of polymerases, and a conditional Poisson sampling model for the generation of site-specific read counts. We recently adapted this model to characterize the equilibrium dynamics of transcription initiation and promoter-proximal pausing based on steady-state NRS data [25], ignoring variability in elongation rate throughout the gene body. Here, we take a complementary approach, focusing on gene-body elongation rates but ignoring promoter-proximal pausing. We focus in particular on the relationships between local rates of elongation and various kinds of genomic and epigenomic features (**Fig. 1**).

**Figure 1:**
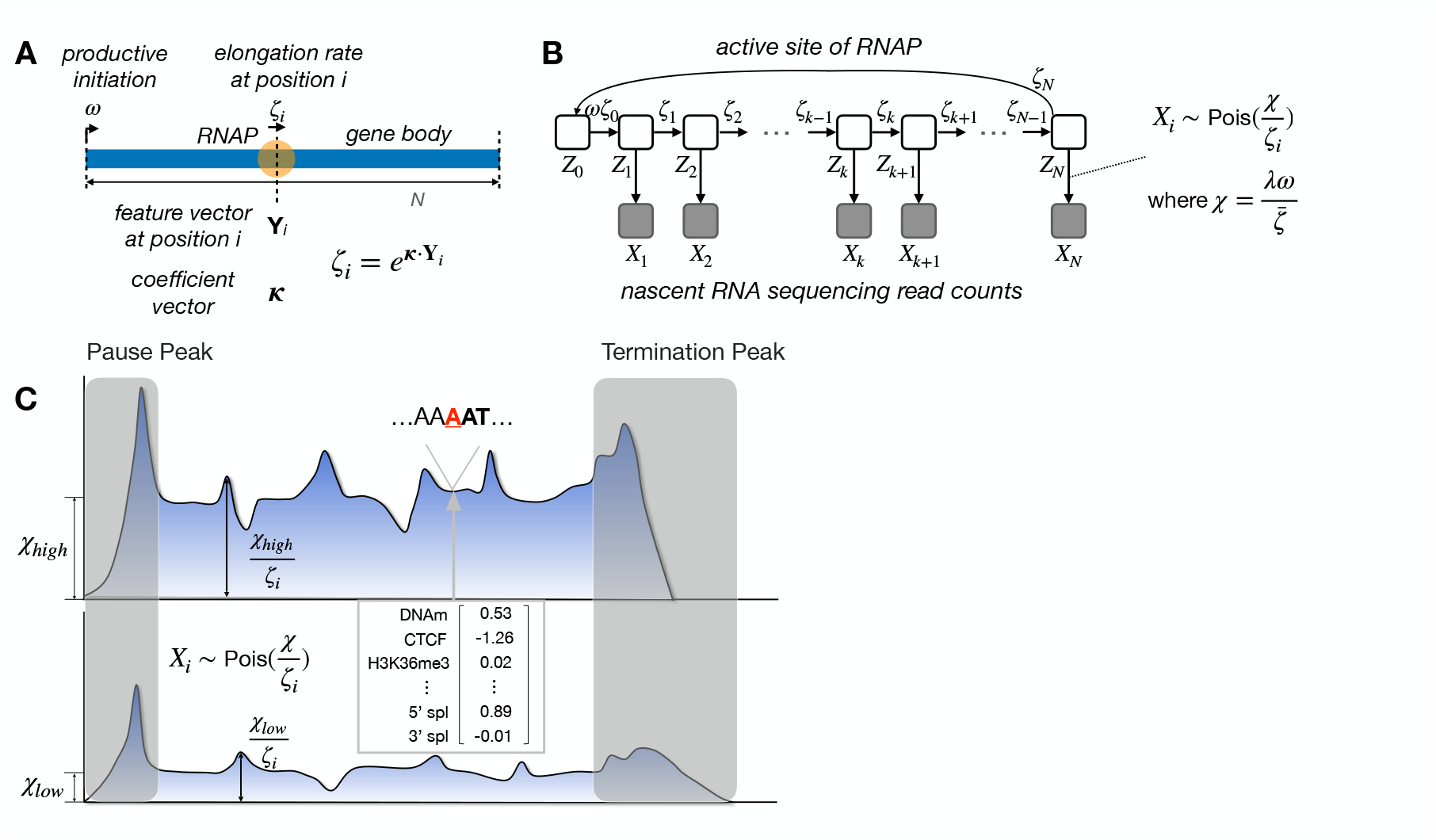
**A**. Conceptual illustration of kinetic model for Pol II movement along DNA template in gene body. At nucleotide site *i*, local elongation rate *ζ*_*i*_ is an exponentiated linear function of features **Y**_*i*_ and coefficients *κ*. Promoter-proximal pausing and termination are ignored here. **B**. Graphical model representation showing unobserved continuous-time Markov chain (*Z*_*i*_) and observed NRS read counts (*X*_*i*_). **C**. Conceptual illustration showing that differences in average gene-body read depth are explained by the scaled initiation rate *χ*, while relative read depth is explained by the generalized linear model for local elongation rate *ζ*_*i*_. Read count *X*_*i*_ is assumed to be Poisson distributed with mean 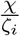. Pause and termination peaks are omitted.

A key challenge in this analysis is that the information about elongation rate at any individual site is weak: it is provided only by local increases or decreases in NRS read-depth, which are often subtle. In addition, the stochastic sampling of Pol II densities via NRS sequencing causes the signal in the raw data to be noisy. We address these problems by describing the relationship between an arbitrary vector of genomic features at each site *i*, denoted **Y**_*i*_, and the elongation rate at that site, *ζ*_*i*_, by a generalized linear relationship, *ζ*_*i*_ = exp(***κ***·**Y**_*i*_), where ***κ*** is a vector of coefficients that is shared across all sites and all genes (**Fig. 1A&B**). In this way, the model can efficiently pool information about local rate across many sites in the genome and circumvent the problem of noise at each site.

A second challenge is that NRS read counts reflect differences in gene-level initiation rates as well as both gene-level and local elongation rates (**Fig. 1C**). In particular, at steady state, the average read depth within each gene body *j* reflects the ratio of the productive initiation rate *ω*_*j*_ to the average elongation rate 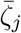 along the gene body [24, 25]; thus, these two influences cannot be disentangled when interpretting differences between genes in average read depth. Therefore, we focus instead on *local* variability in elongation rate. We estimate a separate compound parameter *χ*_*j*_ for each gene *j*, which can be interpreted as a read-depth-scaled initiation-to-elongation rate ratio for the gene as a whole (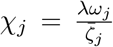; see **Methods**). Then we allow each site *i* in gene *j* to have a different *relative* local elongation rate *ζ*_*i,j*_, which is defined as a function of the local features via the generalized linear model. These *ζ*_*i,j*_ can be thought of as scale factors for the average elongation rate, taking values *<*1 for local slow-downs and values *>*1 for local speed-ups in polymerase movement. The model accounts for the observed read-depth *X*_*i,j*_ at each site *i* by assuming *X*_*i,j*_ is Poisson-distributed with mean *χ*_*i*_*/ζ*_*ij*_. Thus, the model accounts for genewise differences in average read depth using the free *χ*_*j*_ parameters, but explains local variation within each gene using the generalized linear model.

### The model recovers true elongation rates and epigenetic correlates in realistic simulations

We first tested our modeling approach on simulated data, making use of a recently developed simulator for nascent RNA sequencing data called SimPol (“Simulator of Polymerases”) [25]. SimPol works by tracking the movement of individual polymerases along the DNA templates in thousands of cells, under user-defined initiation, pause-escape, and elongation rates. After the Pol II density along the DNA templates reaches equilibrium, the program uses Poisson sampling to generate synthetic NRS read counts that reflect that density. These read counts are designed to be realistically sparse. For a typical gene in our simulations, the majority of nucleotides have counts of zero, most of the remainder have counts of one, and only a few have counts of greater than one (**Fig. 2A**; **Methods**). Importantly, SimPol tracks potential collisions between polymerases and prohibits one polymerase from passing another along the same template, despite that our model ignores these phenomena.

**Figure 2:**
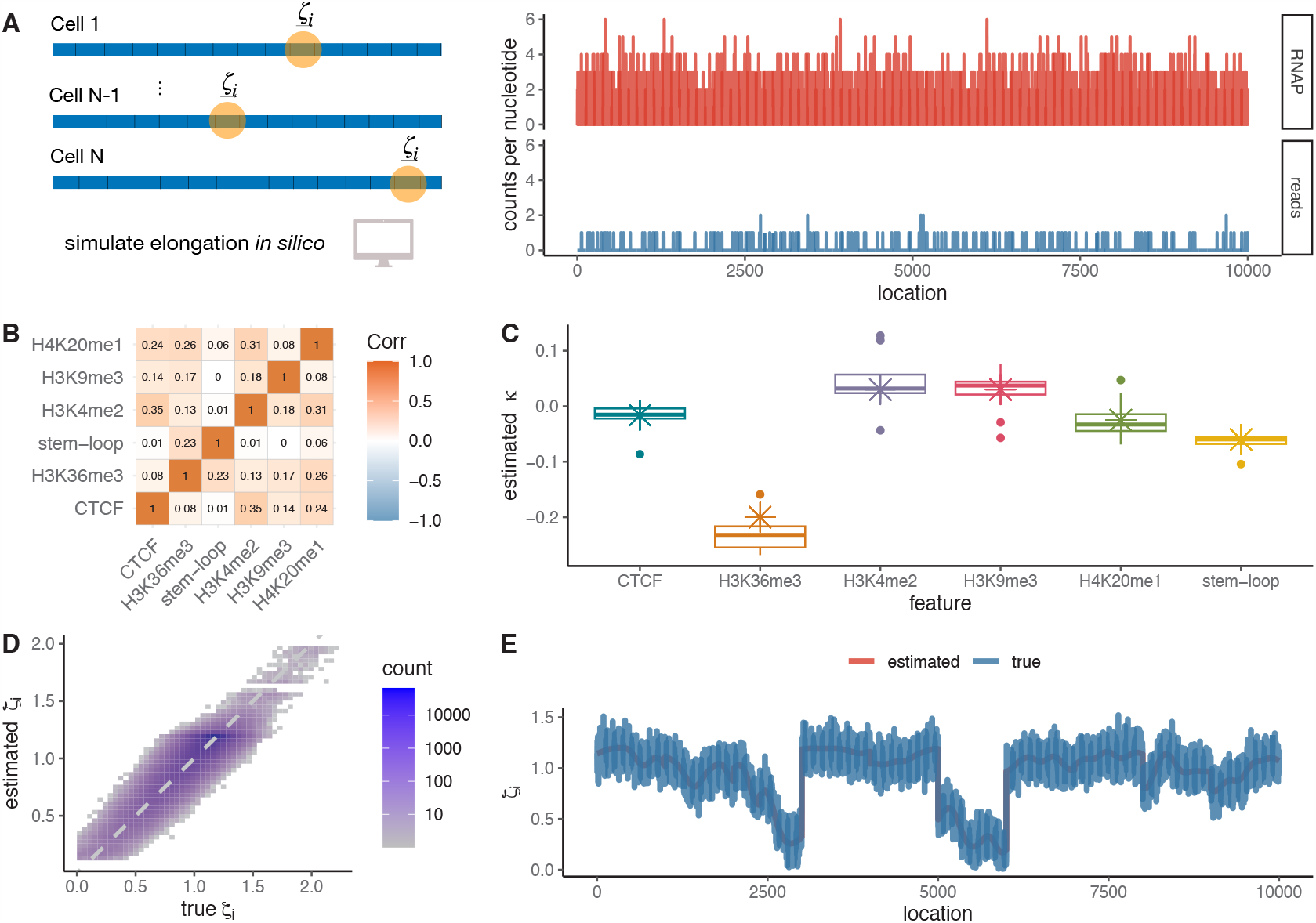
**A**. The SimPol simulator tracks the movement of virtual polymerases across DNA templates in a population of cells (*left*). Once an equilibrium is reached, read counts per site are sampled in proportion to the simulated Pol II density, such that the average read depth is matched to real PRO-seq data (*right*). **B**. Correlation map of selected epigenomic features for simulations (Spearman’s *ρ*). **C**. Box plots for estimated coefficients *κ* in ten replicates compared with ground truth in simulations (crosses). **D**. Estimated vs. true nucleotide-specific elongation rates *ζ*_*i*_ across all simulated TUs (*r*^2^ = 0.748). **E**. Estimated vs. true nucleotide-specific elongation rates *ζ*_*i*_ along an individual TU in ten replicates (*r*^2^ = 0.869).

For this study, we ignored the components of SimPol concerned with promoter-proximal pausing but extended the simulation scheme to allow for correlations between the elongation rate at each nucleotide site and a variety of (synthetic) epigenomic covariates. We based our synthetic covariates on real data from K562 cells (as discussed further below), focusing on CTCF transcription binding sites, four different histone marks, and RNA stem-loops as representative features. We tiled each synthetic DNA template with associated covariates using a block-sampling approach based on real data, to maintain realistic genomic densities and correlation patterns among the covariates (**Fig. 2B**; see **Methods**). We then determined the “true” elongation rate at each site as the sum of an exponentiated linear function of the associated covariates (using coefficients derived from real data) and independent Gaussian “noise,” to simulate other sources of variation not included in the model. These true per-nucleotide elongation rates were passed to SimPol such that the same per-nucleotide elongation rates applied to each of the 5,000 cells used in data generation. While our simulation scheme shares some assumptions of our model, it is nevertheless useful for evaluating our estimation of both the GLM coefficients and the predicted local elongation rates from sparse NRS data in the presence of correlated covariates.

We carried out 10 identical experimental replicates with SimPol, each time simulating NRS data for 100 transcription units (TUs) 10 kbp in length. Each TU had its own initiation rate, sampled from estimates from real data (see **Methods** and **Supplementary Fig. S1A**). For each replicate, we used the simulated NRS data for 80 of the simulated TUs as “training” data (for parameter estimation), and set aside the remaining 20 as “testing” data. We then estimated the coefficient vector ***κ*** from the training data by maximum likelihood under our model (see **Methods**). Overall, the estimated coefficients showed excellent agreement with the true values, with some variation across replicates (**Fig. 2C**). As expected, some of the sparser covariates— such as H3K4me2—resulted in somewhat greater variance in the estimated coefficients than did the denser covariates—such as the stem-loops. We observed a slight estimation bias in some cases (most notably, H3K36me3), perhaps owing to unmodeled correlations between covariates. In most cases, however, the estimated coefficients appeared to be approximately unbiased, with median values close to the truth.

In a second experiment, we re-estimated the coefficient vector ***κ*** from all “training” TUs, pooled across replicates (10 replicates × 80 TUs = 800 TUs), and used these estimates to predict per-nucleotide values of the elongation rate *ζ*_*i*_ in each of the 20 × 10 = 200 TUs held out for testing. In this setting, where the “true” values also reflect a generalized linear model, the predicted elongation rates were well correlated with the true values (*r*^2^ = 0.748; **Fig. 2D**), with the unexplained variance approximately equal to the contribution of Gaussian noise to the true values (∼25%; see **Supplementary Fig. S2**). A version without the addition of Gaussian noise showed almost perfect performance (**Supplementary Fig. S2A–D**). When visualized along an individual TU, the predictions can be seen to form a smooth line near the middle of the cloud of site-to-site variability in true rates, which reflects the addition of Gaussian noise (**Fig. 2E**). While the precise degree of predictivity in these experiments depends on the details of our scheme for simulating “true” elongation rates, these results nevertheless demonstrate that our model can produce accurate predictions of *ζ*_*i*_ provided informative covariates are available.

### Several epigenomic and sequence features are correlated with local elongation rate

Having established that our model works well with simulated data, we applied it to real PRO-seq data from K562 cells [26], a cell type for which abundant data is available from epigenomic assays. Based in part on previous reports of features associated with elongation rate [5–8], we selected a diverse collection of twelve features as covariates, including: 5^*/*^ and 3^*/*^ splice sites evident from cell-type-matched RNA-seq data, DNA methylation based on whole-genome bisulfite sequencing data, CTCF binding sites based on ChIP-seq data, six histone modifications (H3K4me1, H3K9me1, H3K9me3, H3K36me3, H4K20me1, and H3K79me2) based on ChIP-seq data (all from ENCODE [27]), as well as apparent RNA stem-loops based on DMS data [28] and low-complexity sequences annotated in the UCSC Genome Browser [29]. At this stage, we omitted features strongly correlated with the DNA-sequence base composition (to be addressed in the next section), such as DNA melting temperature and stability of the DNA-RNA duplex. We also excluded several histone marks whose potential association with elongation rate appears to be limited to the region immediately downstream of the TSS, which is not our focus in this study (see **Supplementary Fig. S3**).

Real PRO-seq data has the difficulty that, in addition to reflecting local elongation rates, it is also strongly influenced by phenomena such as promoter-proximal pausing, unannotated transcription start sites (TSSs), overlapping isoforms, and enhancers within TUs. We therefore preprocessed the raw data in several ways (see **Methods** for details). Briefly, we adjusted annotated TSSs as needed using cell-type-matched CoPRO-cap data, which enriches for capped 5^*′*^ ends [30]. We used DENR [31] to select the dominant pre-mRNA isoform for each TU, as supported by our PRO-seq data. We then stringently masked out regions within the selected TUs that could plausibly represent internal TSSs (from enhancers or other TUs), based on dREG predictions [32] and GRO-cap data [33] (see example in **Supplementary Fig. S4**). Because our analysis focuses on elongation dynamics in the gene body, we also eliminated the first 2250 bp downstream of the TSS—which appeared to include all promoter-proximal pause sites—as well as the final 250 kbp upstream of the annotated transcription termination site (TTS).

Following the application of these filters, the remaining PRO-seq read depths still exhibited a characteristic “U-shape” along gene bodies (**Supplementary Fig. S5**), which has previously been noted and hypothesized to reflect a tendency for acceleration of Pol II following pause-release and deceleration prior to termination [6]. To ensure that this broad pattern did not interfere with our more local analysis, we followed previous work [31] in adjusting the read depths so that they were globally uniform on average. Finally, because our features of interest have quite different levels of precision along the genome—e.g., splice-site annotations pinpoint individual nucleotides whereas ChIP-seq-based signals have resolutions of hundreds or thousands of nucleotides—and because features can influence elongation rate at adjacent nucleotides, we devised “smoothing filters” that could be applied to each feature, distributing its information along the genome sequence in an appropriate manner. This system essentially allows the features to be placed on the same genomic scale as one another and as our nucleotide-level PRO-seq data (see **Methods** and **Supplementary Fig. S6A–C**). After filtering, we evaluated the correlation structure of the epigenomic features and found relative low correlation overall, except among the histone marks where correlation was stronger (**Supplementary Fig. S6D**).

We applied our model to the filtered data for a large collection of robustly expressed protein-coding genes, first selecting the 6,000 genes with the highest PRO-seq signals in their gene bodies, then randomly sampling 2,000 genes in each of ten rounds of analysis. For each sample of 2,000 genes, we estimated the coefficient vector *κ* by maximum likelihood, and we used the variation in these estimates to obtain standard errors for the coefficients. We found that the estimated coefficients were mostly negative in sign, indicating associations with a reduction in elongation rate (**Fig. 3A**). The strongest signal, by far, was associated with DNA methylation (*κ* = −0.20). Moderately strong reductions in elongation rate were also associated with H3K36me3 (*κ* = −0.095), H3K9me1 (*κ* = −0.067), and RNA stem-loops (*κ* = −0.051). Somewhat weaker reductions were associated with both 3^*/*^ (*κ* = −0.033) and 5^*/*^ (*κ* = −0.019) splice sites, CTCF binding sites (*κ* = −0.022), and other histone modifications (H4K20me1, H3K9me3, H3K4me1; (*κ* ∈ [−0.041, −0.024]). Only two coefficients were positive, indicating associations with increases in elongation rate: those for low-complexity sequence (*κ* = 0.016) and the H3K79me2 histone mark (*κ* = 0.030).

**Figure 3:**
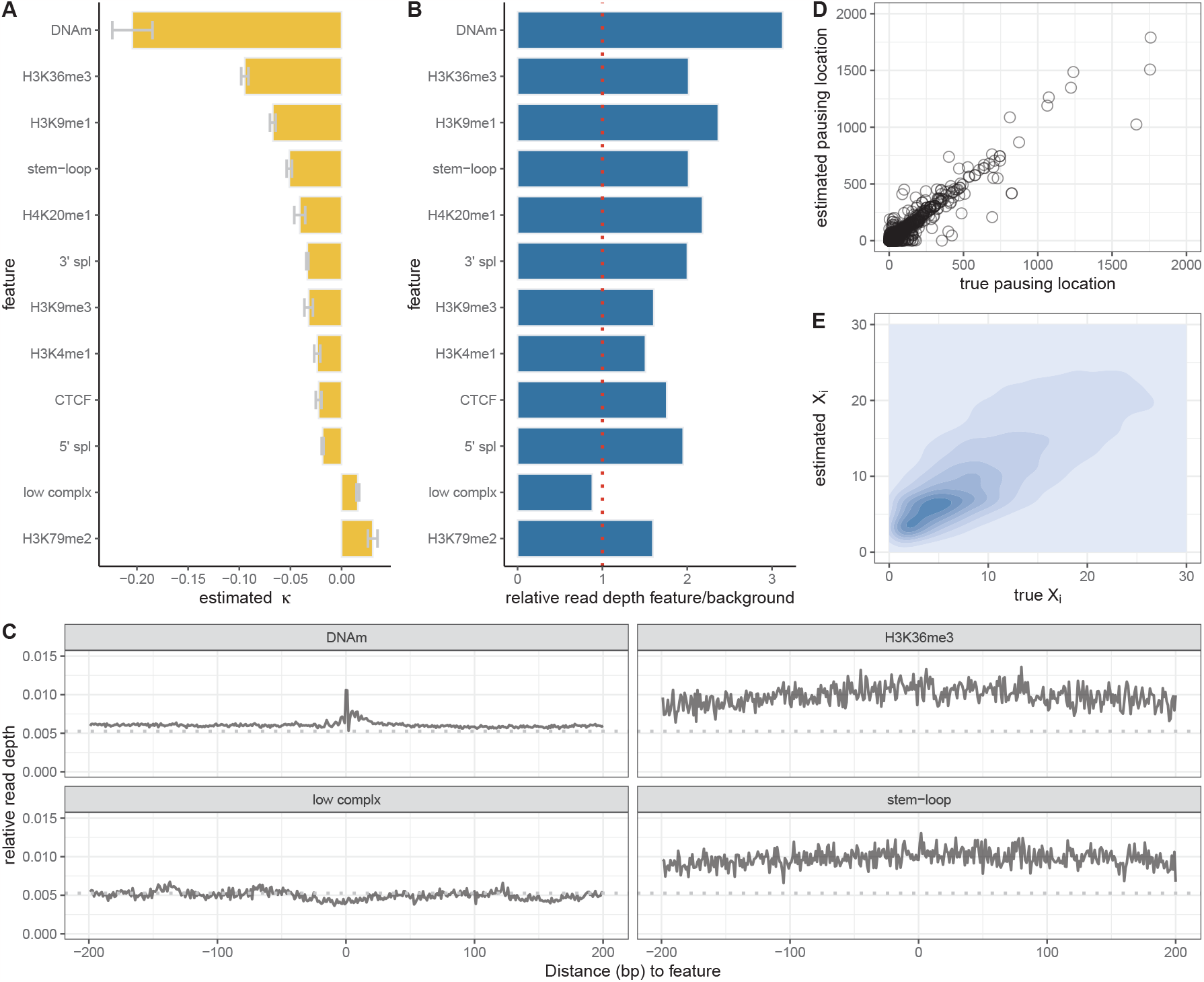
**A**. Estimated coefficients *κ* for the twelve epigenomic features considered, based on PRO-seq data for K562 cells [26]. Sign indicates direction and absolute value indicates strength of correlation with local elongation rate. Error bars indicate one standard error in each direction. **B**. Ratio of relative average PRO-seq read depth in regions covered by each feature to that in regions not covered by it (see text). **C**. Metaplot of relative read depths centered on four selected features. Dashed line represents average across all gene bodies. **D**. Estimated vs. true locations of pausing locations within gene bodies (see text) (*r*^2^ = 0.60). **E**. Predicted vs. true PRO-seq read depths (*X*_*i*_) for held-out data averaged over 1kb intervals for all TUs (*r*^2^ = 0.28).

These observations are generally consistent with previous observations at the level of entire genes, with negative correlations having been reported for DNA methylation [6, 7], H3K36me3 [6], H4K20me1 [4], splice sites [17, 22, 34], and CTCF binding [17, 19, 21], and positive correlations having been reported for H3K79me2 [6–8] and low-complexity sequences [7]. Our analysis shows, however, that changes in the presence or absence of these features are directly associated with *nearby* changes in elongation rate, further suggesting underlying mechanistic relationships with the movement of Pol II (see **Discussion**).

To validate that our model-based associations were well supported by the raw data, we carried out a simpler, non-model-based analysis that compared the average read depths in sites within gene bodies that were annotated (“covered”) and were not annotated (“noncovered”) by each feature (**Fig. 3B**). To account for differences in initiation and/or average elongation rates, we first divided all read counts by their average value within each gene body. We then examined the ratio of the resulting relative read depth in the “covered” regions to that in the “uncovered” regions, pooling data across all 6,000 robustly expressed genes. A ratio of *>*1 therefore indicates an increase in PRO-seq read depth at annotated sites, whereas a ratio of *<*1 indicates a decrease in read depth.

This simple measure was generally consistent with the estimated coefficients under our model, with features that had negative coefficients under the model (indicating an association with reduced elongation rate) showing ratios *>* 1 and features that had positive coefficients showing ratios *<* 1 (**Fig. 3B**). The only exception was the H3K79me2 histone modification, which showed a weakly positive estimate of *κ*. but a slight enrichment for higher read PRO-seq read counts. This difference may be attributable to coincidence of H3K79me2 marks with other features, which is not considered in the non-model-based analysis. The tendency for a local increase or decrease in read depth associated with genomic annotations could also be clearly observed in metaplots centered on annotated sites (**Fig. 3C**). Notably, among the features with negative coefficients, the absolute values of the coefficients were also roughly consistent with the fold-changes in read depth (**Fig. 3B**), with some differences in rank order probably owing to the independent consideration of each feature in the non-model-based analysis. Overall, this comparison shows that our model-based analysis does faithfully reflect first-order patterns of relative read depth, but makes a number of adjustments in magnitude—and occasionally in sign—by jointly considering all features together in one unified framework.

Another way to validate our model would be to test its ability to predict held-out PRO-seq data. However, the predictive power for read counts at individual nucleotide sites is poor, even with simulated data, because the data are so sparse (see **Supplementary Fig. S2D&E**). We carried out two modified experiments to circumvent the problem of sparse data. First, we tested the ability of the model to predict “pausing locations” within gene bodies, defined as 200-bp intervals having the highest average read counts for each gene (see **Methods**). We found, even with held-out genes, that model-based predictions of such pausing locations agreed quite well with the truth (*r*^2^ = 0.60; **Fig. 3D**). Second, we tested the ability of the model to predict average read counts across larger windows, spanning 1000 bp. At this level of resolution, the predictions for held-out genes are still approximate but much better than for individual nucleotides (*r*^2^ = 0.28; **Fig. 3E**). These results suggest that the model is effective at getting at underlying elongation rates even if the predictive power for nucleotide-level PRO-seq read counts remains weak.

### An extension to the model accommodates DNA-sequence *k*-mers

In addition to epigenomic factors, it is well known that DNA sequences can also influence local elongation rates, based on studies ranging from bacteria [35] to yeast [36] and mammals [37] (reviewed in [14]). Promoter-proximal pausing in *Drosophila* and mammals is also associated with particular sequence motifs [9, 37–39]. Recently it was shown using NET-seq and PRO-seq data for *S. cereviseae* that nucleotide 5-mers were strongly predictive of local elongation rates, beyond what could be explained by G+C content, DNA folding energy, or sequencing bias [11].

We therefore extended our generalized linear model to consider the *k*-mer content of the local DNA sequence. Following ref. [11], we initially considered 5-mers only (*k* = 5). These 5-mers are accommodated using indicator features in our regression framework: at each position *i*, the indicator feature associated with the 5-mer centered at *i* is set to 1 and the remaining 5-mer indicator features are set to 0. These features can be used alone or together with epigenomic features.

The challenge with this strategy is that it requires a high-dimensional feature vector. The 5-mer features alone number 4^5^ = 1024. We therefore added a sparsity penalty to our likelihood function, which has the effect of limiting the set of features that have non-zero coefficients and forcing the model to choose the ones that are most informative. After experimentation, we settled on an L1 (lasso) penalty and determined the strength of the penalty by cross-validation (see **Methods** and **Supplementary Fig. S7** for details).

We first tested this approach with simulated data. Briefly, we sampled 20,000 sequences of length 1 kbp from the human genome. We randomly drew 100 5-mers from these sequences and assigned them negative or positive correlations with local elongation rate, with *κ* values ranging from −0.3 to +0.3. All other 5-mers were assigned *κ* = 0, indicating no correlation. Based on these coefficents, we then generated “true” nucleotide-specific elongation rates along the sampled DNA sequences under the assumptions of our exponentiated linear model. As in our previous experiments, we passed these local elongation rates to SimPol, which used them to generate synthetic NRS read counts. When we fitted our GLM to these sequences, using 5-mer features only and the L1 penalty, non-zero coefficients were assigned to 100 features, which heavily overlapped the set of features that had truly been assigned non-zero coefficients (**Supplementary Fig. S7**). In addition, the estimates of *κ* were generally close to the true values (**Fig. 4A**; *r*^2^ = 0.89) as were the predictions of local elongation rate (**Supplementary Fig. S7D**; *r*^2^ = 0.88). Overall, the method appears to be effective both at recovering 5-mers correlated with elongation rate and at predicting the local elongation rate itself.

**Figure 4:**
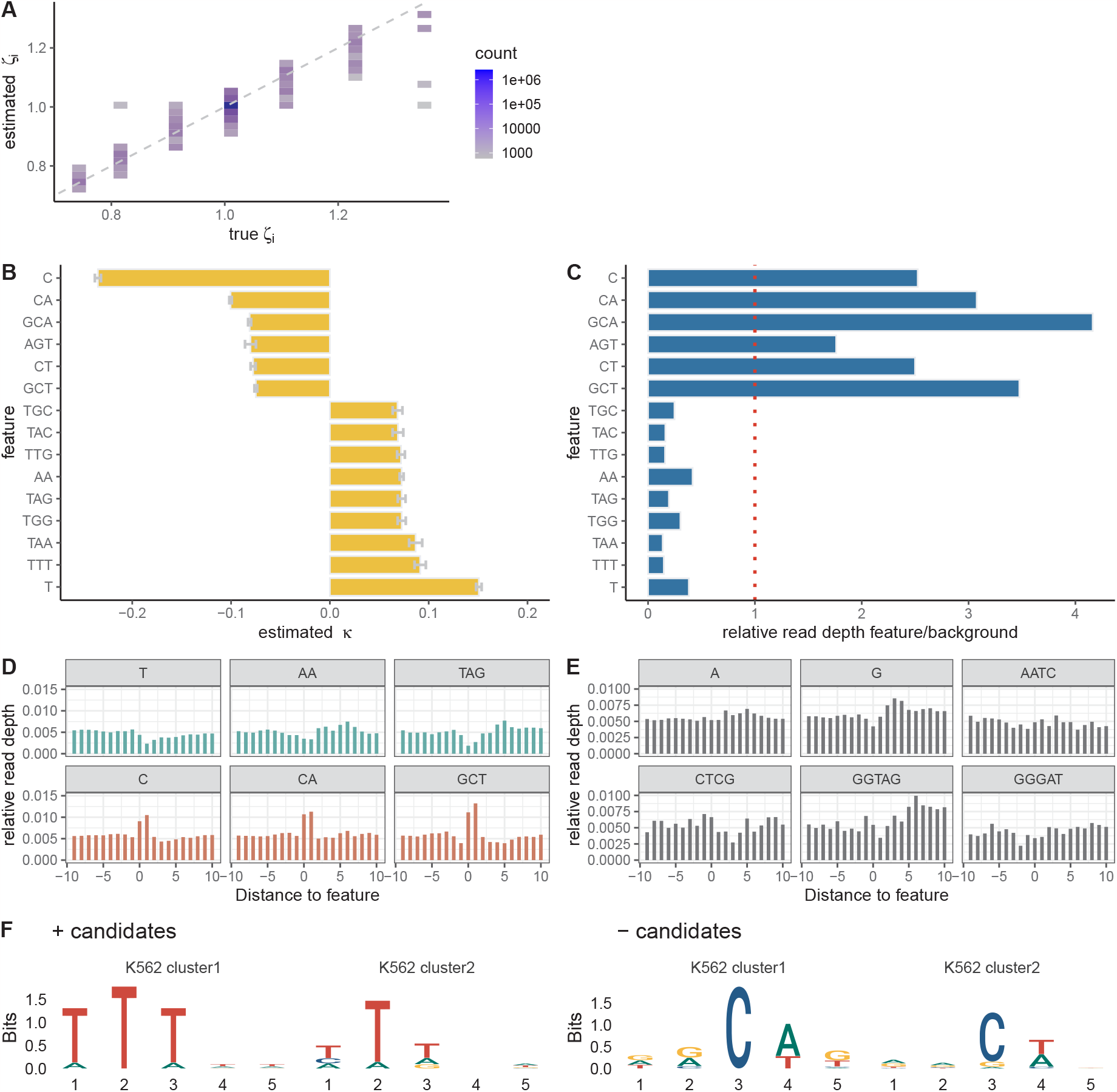
**A**. Estimated vs. true nucleotide-specific elongation rates *ζ*_*i*_ in ten rounds of simulated *k*-mer data (*r*^2^ = 0.89). **B**. Estimated coefficients *κ* for top *k*-mers (*k* ≤ 5) based on PRO-seq data for K562 cells [26]. Sign indicates direction and absolute value indicates strength of correlation with local elongation rate. Error bars indicate one standard error in each direction. **C**. Ratio of relative average PRO-seq read depth at sites associated with each *k*-mer to that at sites not associated with it (see text). **D**. Metaplot of relative read depths for three *k*-mers with positive coefficients (*top*) and three with negative coefficients (*bottom*). **E**. Metaplot of relative read depths for six *k*-mers having coefficients close to zero. **F**. Sequence logos summarizing clusters of 5-mers five nucleotides centered on the active site that are positively (*left*) or negatively (*right*) associated with elongation rate.

### Several *k*-mers are strongly associated with local elongation rates in K562 cells

With this sparse-regression, *k*-mer-based version of the model in hand, we re-analyzed the PRO-seq data from K562 cells, this time considering DNA sequence *k*-mers only (i.e., omitting the epigenomic features; they will be re-introduced below). We started with the same 6,000 genes as in the epigenomic analysis but this time sampled four batches of 500 genes to limit computational cost. Also, instead of considering 5-mers only, we allowed for *k*-mers of any length up to and including five nucleotides (*k* ∈ {1, 2, 3, 4, 5}). In this version of the model, the *k*-mers of different sizes compete with one another, and the smallest *k*-mer that adequately explains the data will tend to be selected. For example, if the correlations with elongation rate are truly driven by G+C content, the model will tend to choose the G and C 1-mers rather than many different 5-mers containing Gs and Cs. In addition, because the model is additive, larger *k*-mers are assigned coefficients representing their contributions *beyond* those of shorter *k*-mers nested within them. For example, if both G and AG are included in the model, then the coefficient assigned to G will reflect its marginal contribution and the coefficient assigned to AG will reflect only the additional contribution of a preceding A.

When we fitted the model to the K562 PRO-seq data we found that the strongest signals, by far, were for a negative correlation of cytosine (C) nucleotides (*κ* = −0.24) and a positive correlation of thymine (T) nucleotides (*κ* = 0.15) with local elongation rate (**Fig. 4B**). The negative correlation was enhanced when the C was followed by an A (CA) or a T (CT) or when it was preceded by a G (GCA, GCT). The nucleotide trimer AGT was also negatively correlated with elongation rate. The positive correlation of a T was enhanced when it was preceded or followed by additional Ts (TTT, TTG). In some cases, a positive correlation also occurred with As or Gs in place of Ts in the central position (TAA, TAG, AA, TAC, TGG, TGC). In general, the *k*-mers associated with increased elongation rate were A+T-rich, with some presence of Gs, and they were particularly enriched for A/T dinucleotides (TT, TA, AA). The negative correlation with cytosines echoes similar findings in *E. coli* [35] and findings for promoter-proximal pausing in mammals [9, 37, 39], and the positive correlation with A+T-rich sequences is consistent with reports of negative correlations with G+C content [6, 7, 15] with some differences (see **Discussion**). Altogether, between 159 and 232 *k*-mers were assigned non-zero coefficients across replicates (**Supplementary Fig. S8B**).

As with the epigenomic version of the model, we validated these *k*-mer associations by examining the corresponding relative read depths. We found that the *k*-mers that had negative coefficients (implying slower elongation rates) did indeed exhibit higher relative read depths, and the *k*-mers associated with positive coefficients (implying faster elongation rates) did have lower relative read depths (**Fig. 4C**). In this analysis, the larger *k*-mers had more divergent relative read depths despite having smaller absolute *κ* estimates, because, as noted, the *κ* estimates reflect only the additional contribution associated with the larger *k*-mer context. For example, GCA has higher relative read depth than CA, which in turn has a higher relative read depth than C, but the *κ* estimates for GCA and CA are smaller than for C because they represent only the additional contributions. A trend in the reverse direction can be seen with T and TTT. When relative read depths for each *k*-mer are plotted along the genome sequence, the local departures from the background levels can be clearly observed (**Fig. 4D**). By contrast, the relative read depths for insignificant *k*-mers are much less pronounced (**Fig. 4E**).

For comparison with the *k*-mers of various sizes, we also analyzed the data with a version of the model that allowed for 5-mers only. To make sense of the identified *k*-mers, we separately clustered the ones positively or negatively associated with elongation rate and summarized each cluster using a sequence logo (**Fig. 4F**; see **Methods** for details). The clusters negatively associated with elongation rate were clearly dominated by a central C, which tended to be followed by A or T and tended to be preceded by G or A. The positively associated clusters were clearly dominated by Ts with a secondary signal from As.

We also evaluated the performance of the *k*-mer model in predicting pausing locations within gene bodies and held-out read counts in 1 kbp intervals (**Supplementary Fig. S9A&B**). While it performed slightly better at predicting pausing locations (*r*^2^ = 0.64 compared with *r*^2^ = 0.60 for the epigenomic model), its predictions of read counts were substantially better (*r*^2^ = 0.65 compared with *r*^2^ = 0.28).

A possible concern with this analysis is that the *k*-mer associations we find with elongation rate, which reflect nucleotide preferences at the 3^*′*^ ends of aligned PRO-seq reads, might be influenced by biases in the PRO-seq protocol. Several lines of evidence, however, suggest that such biases are not driving our results, including comparisons of the bulk distribution of 3^*′*^ nucleotides across data sets (**Supplementary Fig. S10A&B**), an analysis of possible ligation biases (**Supplementary Fig. S10C**,**D&E**), and analyses with a version of the model that explicitly allows for 3^*′*^ nucleotide biases (**Methods** and **Supplementary Fig. S9C**). It is also notable that a strong preference for cytosines at sites of promoter-proximal pausing has been noted with NET-seq data as well [9], and the NET-seq protocol does not include the biotin run-on step of PRO-seq (see [30]). Overall, while we cannot rule out some influence from the protocol on our *k*-mer associations, these findings suggest that any such bias should be minimal (see **Discussion**).

### Most associations with local elongation rate are shared across cell types

To examine the generality of these correlations with local elongation rate, we carried out a similar analysis of PRO-seq data sets from three other cell types: CD14+ [31], MCF-7 [40] and HeLa-S3 [41] cells. These data sets were generated by three different laboratories using slightly different methods, making them also informative about the sensitivity of our conclusions to variations on the PRO-seq protocol. For comparison with our results for K562 cells, we analyzed them separately with the epigenomic and *k*-mer versions of our model. Epigenomic data was available from the ENCODE project for all three cell types [27].

We found that the epigenomic correlates were generally fairly consistent across cell types overall, with the exception of DNA methylation, which showed a strong negative correlation with local elongation rate in K562 and MCF-7 cells, but a strong positive correlation in CD14+ and HeLa-S3 cells (**Fig. 5A**). On further inspection, we found that this difference traced back to secondary TSSs within TUs (e.g., from transcribed enhancers), which were filtered out in K562 and MCF-7 cells using available GRO-cap data but could not be efficiently identified and removed in CD14+ and HeLa-S3 cells, for which no GRO-cap or PRO-cap data was available. An analysis of K562 cells with and without this filter for internal TSSs shows that the correlation with DNA methylation is highly sensitive to the filter (**Supplementary Fig. S11**). Because these internal TSSs tend to be both unmethylated and transcribed, leading to a high PRO-seq read depth, they appear in the context of our model to suggest that unmethylated DNA has a reduced elongation rate, implying a positive correlation between methylation and elongation rate. This spurious signal of positive correlation overwhelms the true signal for negative correlation that occurs elsewhere throughout the gene bodies. We therefore disregard the positive *κ* estimates for CD14+ and HeLa-S3 cells as artifacts of unfiltered internal TSSs. Once DNA methylation was excluded, the *κ* estimates for K562, CD14+, and HeLa-S3 were all strongly correlated, with pairwise *r >* 0.85 in all cases (**Supplementary Fig. S12A**). The MCF-7 data set showed somewhat weaker correlation with the others (*r* ≈ 0.7 with K562 and HeLa-S3, *r* = 0.43 with CD14+).

**Figure 5:**
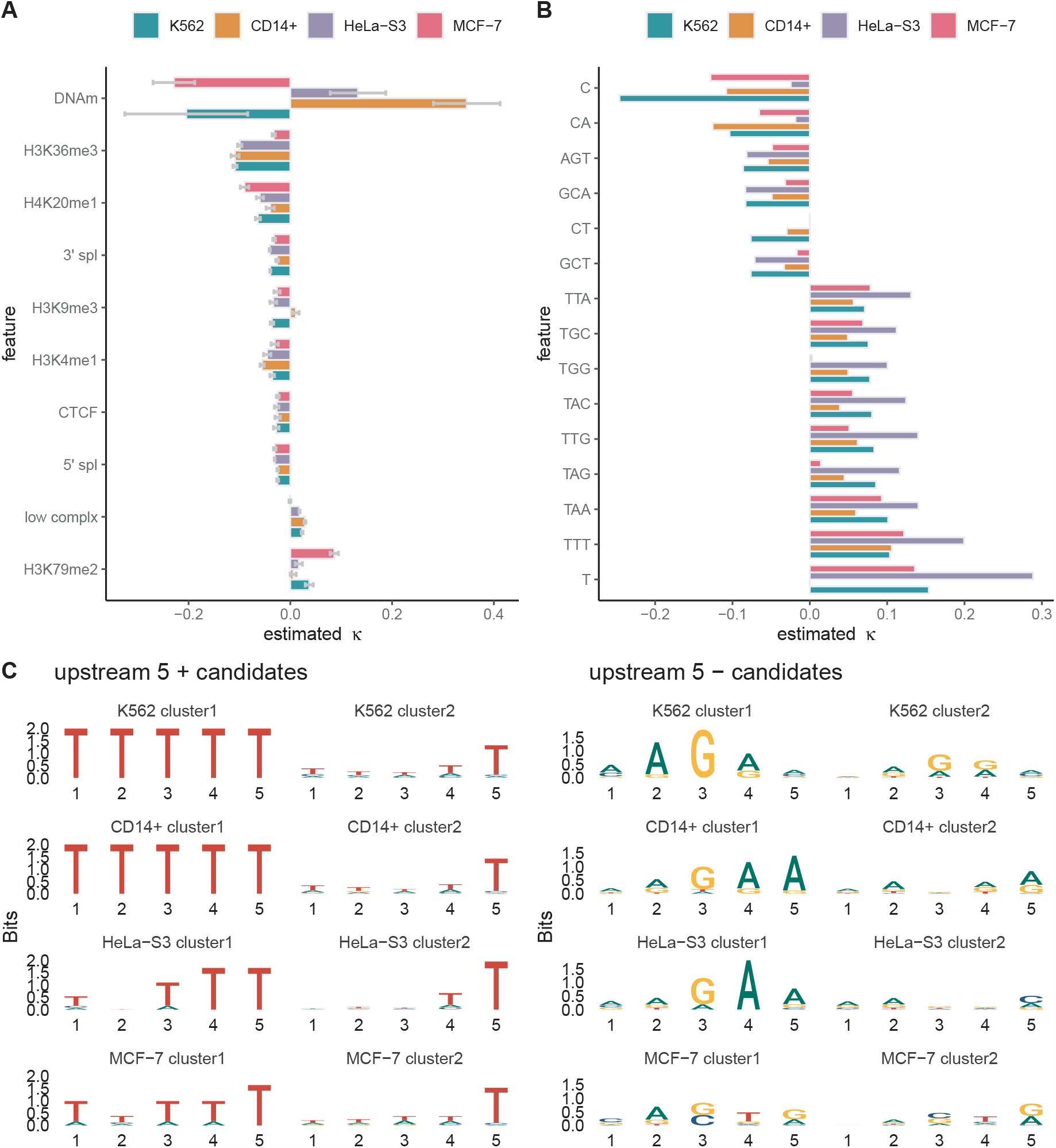
**A**. Estimated coefficients *κ* for twelve epigenomic features based on PRO-seq data for four mammalian cell lines: K562 [26], CD14+ [31], HeLa-S3 [41], and MCF-7 [40]. Sign indicates direction and absolute value indicates strength of correlation with local elongation rate. Error bars indicate one standard error in each direction. **B**. Estimated coefficients *κ* for top *k*-mers (*k* ≤ 5) in the same cell lines. **C**. Sequence logos summarizing clusters of 5-mers five nucleotides upstream of the active site that are positively (*left*) or negatively (*right*) associated with elongation rate (see **Methods**).

We observed a few other, more minor, differences in epigenomic correlations. For example, the positive correlation of H3K79me2 with local elongation rate was considerably strengthened in MCF-7 cells, whereas the positive correlation of low-complexity sequences was lost in MCF-7. These differences traced back to an anomalous pattern of relative read depths around these two features in the MCF-7 data, with increases in H3K79me2 and decreases in low-complexity sequences relative to the background, for reasons we could not discern. Conversely, the negative coefficient for H3K36me3 was smaller in MCF-7 cells. These differences contributed to the reduced global correlation between MCF-7 and the other cell lines—particularly with CD14+, which itself was an outlier in having a positive H3K9me3 coefficient.

With the *k*-mer version of the model, we also observed general agreement across cell types, with pairwise *r* values of ∼0.8 or greater in all cases (**Supplementary Fig. S12B**). Among the most prominent *k*-mers (**Fig. 5B**), the most striking difference was in the *κ* estimate for cytosines, which in CD14+ and MCF-7 cells was about half that in K562 cells, and in HeLa-S3 cells was nearly cut to zero. On further investigation, we found that the HeLa-S3 data set has a different bulk distribution of 3^*′*^ nucleotides, with much less enrichment for cytosines than the others (**Supplementary Fig. S13**). This difference may also help to explain the unusually strong positive correlation for T-containing *k*-mers in HeLa-S3 cells. Despite reprocessing the raw data and re-mapping the reads for HeLa-S3, we were unable to uncover the reasons for this difference. One possible contributing factor, however, is pronounced aneuploidy in HeLa-S3 cells, which can distort copy numbers of transcripts and create challenges in read mapping.

The estimates for cytosines in CD14+ and MCF-7, while reduced compared with K562, were still among the largest (in absolute value) *κ* estimates for those cell types, indicating that cytosines are associated with a substantial reduction in elongation rate across cell types. Nevertheless, these differences in absolute value do suggest that the strength of the correlation measured by our model may be sensitive to details of the PRO-seq protocol. Notably, the CD14+ data set is based on a 2 dNTP run-on protocol (with UTP and CTP), whereas the others are based on 4 dNTP run-on protocols.

### Some distinct *k*-mers are associated with elongation rate upstream and downstream of the active site

We also wondered whether *k*-mers upstream or downstream of the active site might be correlated with elongation rate, for example, owing to the energetics of the DNA-RNA hybrid, Pol II-nucleic acid interactions, or structure in the nascent RNA. We therefore applied the *k*-mer model to the data sets for all four cell types, but this time, instead of considering *k*-mers centered at the 3^*′*^ end of aligned PRO-seq reads, we separately considered *k*-mers that were shifted upstream (in the 5^*′*^ direction) or shifted downstream (past the nascent RNA) by 5, 10, 15, 20, or 25 nt. For this analysis, we used the 5-mer-only version of the model, followed by clustering of 5-mers positively or negatively associated with elongation rate, as introduced above (see also **Methods**).

Upstream of the active site, we found that the *k*-mers positively correlated with elongation rate were dominated by A+T-rich sequences, somewhat like those at the active site itself. Farther upstream, however, they included As and Ts in roughly equal proportions, whereas closer to the active site, they were clearly dominated by Ts (**Fig. 5C**; **Supplementary Fig. S14**). By contrast, the upstream *k*-mers negatively correlated with elongation rate showed a distinctive enrichment for A and G, particularly in alternating patterns. These patterns are reminiscent of GAGA-factor binding sites, which are known to be associated with promoter-proximal pausing of Pol II [34, 38, 42]. These patterns were most pronounced at −5 nt, weaker at −10 nt, and no longer evident by −15 nt, where they were replaced by weakly C+T-rich sequences. Notably, the alternating A and G pattern was not strongly evident at the active site itself, although one of the identified *k*-mers was consistent with it (AGT; see **Fig. 4B**).

Downstream of the active site, both the positively and negatively correlated *k*-mers were much less well-defined. The *k*-mers with positive coefficients, however, were still somewhat A+T-rich, with a preference for T over A. The *k*-mers with negative coefficients at +5 nt still showed some preference for A and G, but it was much less pronounced than upstream of the active site (**Fig. 5C**; **Supplementary Fig. S15**).

### Predicted local elongation rates are available as UCSC Genome Browser tracks

The separate epigenomic and *k*-mer versions of the model both exhibited reasonably good predictive performance on held-out PRO-seq data (**Fig. 3D&E** and **Supplementary Fig. S9A&B**), but we wondered if performance could be improved by combining all features into one model. We therefore devised a version of the model with all twelve epigenomic features and the *k*-mer features of sizes 1–5, applying the lasso penalty to induce sparsity in the high-dimensional feature vector. We fitted this model to the PRO-seq data for K562 cells and tested it on held-out data, as in the previous experiments. We found that the combined model did perform better than the two separate models, but the improvement relative to the *k*-mer-only model was slight (**Supplementary Fig. S16**). This result suggests that most of the information in the epigenomic model can be extracted from the *k*-mer composition of the underlying DNA sequences, and overall, the *k*-mers are more predictive than the epigenomic features, perhaps owing to their much denser coverage along the genome.

Based on this combined model, we created a UCSC Genome Browser track showing the predicted nucleotide-specific local elongation rates genome wide (**Fig. 6**; available at https://bit.ly/elongation-rate-tracks). This track allow our model-based predictions to be viewed alongside gene annotations, epigenomic data, and many other types of genomic data. In this browser track, each of the four cell types (K562, CD14+, MCF-7, and HeLa-S3) is configured as a separate, selectable subtrack, allowing them to be compared to one another easily. Across much of the genome, the subtracks are similar, but in places, differences occur that can be traced to cell-type-specific epigenomic signatures. These tracks are publicly available either for browsing or for download of raw data.

**Figure 6.**
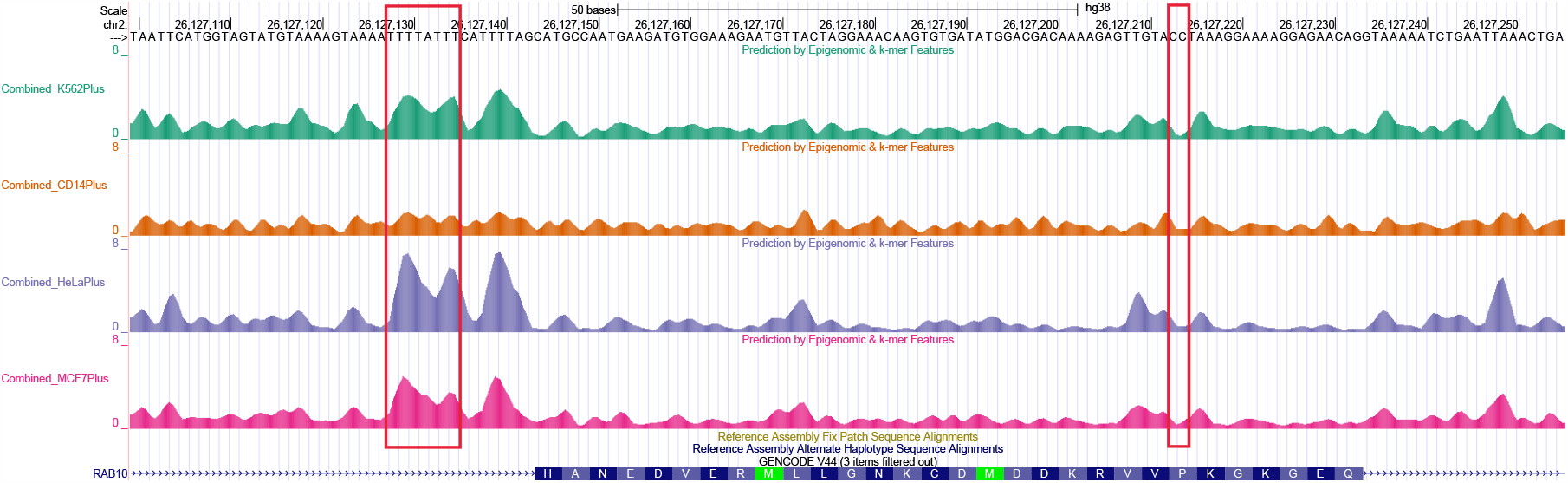
: Screenshot from UCSC Genome Browser track showing predicted local elongation rates for the K562, CD14+, HeLa-S3, and MCF-7 cell types in a region of the *RAB10* gene. These predictions are based on the combined *k*-mer and epigenomic model, but tracks are also available for the epigenomic model only. Notice the elevated predicted rates at poly-T sequences, the reductions at cytosines, and the general reduction throughout the exon.

## Discussion

In this article, we have introduced a new probabilistic model for evaluating correlations between local elongation rate and a wide variety of genomic and epigenomic features. Our model explains nucleotide-specific NRS read counts by assuming they are Poisson distributed with mean inversely proportional to the local elongation rate, which in turn is determined by an exponentiated linear function of nearby genomic and epigenomic features. An optional L1 sparsity penalty can be used to accommodate high-dimensional feature vectors. Simulations show that the model is effective in both identifying correlated features and predicting elongation rates based on those features.

We have separately applied epigenomic and DNA *k*-mer versions of the model to data for four mammalian cell types. DNA methylation emerged as the strongest epigenomic correlate of local elongation rate, followed by H3K36me3 and H3K9me1 histone marks and RNA stem-loops, all of which were associated with slow-downs of Pol II. Other significant negative correlates of rate included splice sites, CTCF binding sites, and several other histone marks. Low complexity sequences and H3K79me2 marks were positively associated with elongation rate. In our DNA *k*-mer analysis, the strongest signals came from cytosines, which were associated with reductions in elongation rate, and thymines, which were associated with increases in elongation rate. We showed that models based on these features have reasonably good prediction power both for pausing locations and held-out PRO-seq read counts. We have used our model to generate publicly available UCSC Genome Browser tracks for the K562, CD14+, MCF-7, and HeLa-S3 cell types.

A key feature of our model is that it directly predicts a *local*, nucleotide-specific elongation rate from features at, or within a few bases of, each nucleotide site. By contrast, most previous studies have focused on correlations at the level of entire genes. The local rate predicted by our model takes values greater than one in regions where Pol II appears to speed up and values less than one in regions where it appears to slow down relative to the average for the gene. Differences in average read-depth betwen genes are separately accommodated through the parameter *χ*, which can be thought of as a gene-wide, read-depth-scaled initiation-to-elongation rate ratio. A disadvantage of our approach is that, because it is based on a single time point and assumes Pol II occupancy is at steady state, it cannot estimate absolute elongation rates. On the other hand, it provides high-resolution preditions of the relative local rate, and it identifies covariates of rate whose presence or absence is physically proximal to changes in NRS read depth. This physical proximity increases the likelihood that these correlations reflect mechanistic, causal relationships—although we still cannot prove causality in this setting.

Despite this difference in our model, the correlations it revealed were generally consistent with previous reports. For example, several studies have identified negative correlations of elongation rate with DNA methylation in mammalian cells [6, 7] (see also [12]). In addition, there have been reports of positive correlations with H3K79me2 [6–8] as well as negative correlations with H3K36me3 [6] and H4K20me1 [4]. It is now well established that elongation rates are negatively correlated with exon density [6, 7, 17], and this relationship appears to be driven, at least in part, by a local reduction in rate at splice sites [17, 22, 34], likely from co-transcriptional splicing. A negative correlation of elongation rate with CTCF binding has also been noted, which in some cases influences splicing [17, 19, 21]. The relationship between RNA secondary structure and elongation rates does not appear to have been examined genome-wide in mammals, but associations of such structure with reduced elongation rates have been observed in bacteria, yeast, and *in vitro* systems [17, 43–45]. A positive correlation of elongation rate with low complexity sequences has also been reported [7]. Our analysis shows that these correlations hold at the local level as well as at the level of averages across genes.

Another important difference is that previous studies have generally considered each potential covariate separately, whereas our model combines them in a unified framework, in such a way that their relative contributions to elongation rate can be compared. As a result, we can say, for example, that the impact of DNA methylation on elongation rate is about twice that of H3K36me3 marks in K562 cells, accounting for all other covariates (**Fig. 3A**). Perhaps owing to our joint model, we do see some minor differences from previous results; for example, H4K20me1 [7] and H3K4me1 [6] have been reported to be positively correlated with elongation rate, whereas we find negative correlations. In an analysis of this kind, the directionality of a relationship can change depending on whether or not other correlated covariates are considered, as was observed with H3K79me2 in **Fig. 3B**.

The negative correlation we observe with a cytosine at the 3^*′*^ end of a nascent transcript echoes a similar finding for promoter-proximal pausing, where paused Pol II shows a strong preference for cytosines at the active site [9, 37, 39]. These studies have also shown some enrichment for G at the preceding position, as in our findings, although the motifs they identified were generally more dominated by Cs and Gs than ours. A similar, but weaker, preference for cytosines was also previously observed outside of the promoter-proximal region based on CoPRO-cap and PRO-seq data [30]. This preference for cytosines has been conjectured to be a consequence of cytosine being the least abundant nucleotide and therefore the slowest to incorporate into nascent transcripts [30]. A similar association between cytosines and pausing of RNA polymerase (RNAP) has been observed in *E. coli*, where it appears to result from RNAP-nucleic acid interactions that inhibit next-nucleotide addition [35] (see also [46]). In this case, however, the preference is for either C or T, both of which tend to be followed by G.

The positive correlation we find between A+T-rich sequences and elongation rate is consistent with many reports indicating a general correlation with G+C content, with slower elongation rates in G+C-rich sequences—and, accordingly, faster rates in A+T-rich sequences—likely resulting from stronger RNA-DNA hybrids and a tendency to form stable RNA secondary structures [6, 7, 15]. It has also been reported that increased elongation rates in A+T-rich introns are stimulated by the U1 snRNP at 5^*′*^ splice sites [47]. Our findings differ from these previous reports, however, in indicating that the effect seems to be driven by thymine somewhat more than adenine bases.

Our analysis assumes that Pol II occupancy along the genome is at a steady-state equilibrium among the cells in the sample. While this assumption undoubtedly does not strictly hold, it seems likely to be reasonably approximated in cell lines that have not been subjected to a treatment or stimulus (control samples). The averaging effect of sequencing a pool of cells should help further in establishing a reasonable proxy for such an equilibrium. Importantly, by operating under this steady-state paradigm, we are able to analyze all expressed genes, not just a subset at which expression can be induced, as in strategies that measure elongation rates from time-course data (e.g., [5–7]). We also avoid the off-target effects of chemical treatments that block initiation or pause escape. Nevertheless, it may be worthwhile, in future work, to extend our generalized linear model for transcription elongation to the nonequilibrium setting and see whether new correlates of elongation rates can be detected from time course data (see [24]).

A second, perhaps more delicate, assumption is that NRS read counts faithfully reflect the density of transcriptionally engaged polymerases across the genome. The main concern here is that read counts could be influenced by biases in library preparation, sequencing, read mapping, or other processes. In principle, any genomic or epigenomic feature favored or disfavored in these processes could spuriously appear to be associated with elongation rate in our analysis. Most of our epigenomic associations should be fairly robust to such biases, since they reflect modifications to the DNA template rather than the nascent RNA. RNA stem loops are a possible exception, but our findings suggest that they are over-represented in PRO-seq read counts, rather than under-represented, as might be expected from interference in capturing structured RNAs.

On the other hand, it is easy to imagine biases that would affect the *k*-mer associations that we detect, particularly the single nucleotides that we find to be over-represented (C) or under-represented (T) at the 3^*′*^ ends of aligned reads. In several follow-up analyses, however, we could find no evidence that biases in the protocol were responsible for the cytosine and thymine associations with elongation rate. First, we found that the bulk distribution of 3^*′*^ nucleotides for PRO-seq reads was fairly consistent across data sets, even when generated by different research groups (**Supplementary Fig. S10A&B**), suggesting that it is not highly sensitive to variations of the protocol. Second, we found that the cytosine enrichment and thymine depletion were present with or without removal of PCR duplicates. Third, we tested directly for a ligation bias, despite that T4 RNA ligase has been reported to have little sequence bias [48]. Specifically, we made use of adapter sets in which the 5^*′*^ adapter incorporates a UMI with randomly occurring nucleotides at its 3^*′*^ end, and compared adapter dimers that contained inserts with ones that did not, finding no difference in their cytosine frequencies and little difference overall (**Supplementary Fig. S10C**,**D&E**. Finally, we devised a “sequence-biased” version of the model that expects the 3^*′*^-most base to appear in proportion to its bulk distribution, under the assumption that an unknown bias drives its frequency, and fitted it to the data, finding that, while the coefficients for C and T were greatly reduced (by design), the coefficients for most larger *k*-mers were relatively unaffected (**Methods** and **Supplementary Fig. S9C**). Altogether, we could find no aspect of the protocol that could explain our associations with C, T, or other *k*-mers. It is also worth noting that NET-seq data—which is produced using a quite different protocol from PRO-seq, without run-on—also shows a strong preference for cytosines at sites of promoter-proximal pausing [9].

To our knowledge, this study represents the first attempt to model local rates of transcription elongation across mammalian genomes. Overall, we find that many features that correlate with elongation rates at the level of entire genes do appear to result in local changes to the elongation rate along gene bodies. By considering these features together in a unified probabilistic model, we can obtain fairly accurate predictions of the local rate, as indicated by our simulations and tests with held-out data. We have made predictions for four cell types publicly available in a UCSC Genome Browser track. We anticipate that they will be useful in a wide variety of downstream analyses, for example, by helping to identify potential cases where transcriptional output is regulated through changes in elongation rate or providing hypotheses about mechanistic influences on elongation rate.

## Materials & Methods

### Unified probabilistic model for nascent RNA sequencing data

Our unified model has been detailed in refs. [24, 25]. Briefly, it consists of two layers: a continuous-time Markov model for the movement of individual polymerases along a TU, and a conditionally independent generating process for the read counts at each nucleotide site (**Fig. 1**). Together, these components produce a full generative model for NRS read counts along the TU, permitting inference of transcriptional rate parameters from the raw data. The model assumes (1) that collisions between polymerases are rare, allowing the movement of each one to be considered independently of the others; and (2) that premature termination of transcription is sufficiently rare that each polyermase can be assumed to traverse the entire DNA template if it is given enough time.

The key extension for the purposes of this work is to allow for a different elongation rate at each nucleotide position, instead of a constant rate across all nucleotides. For reasons that will become clear below, we express the elongation rate for position *i* and gene *j* as a product of a gene-wide average elongation rate 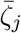 and a position-specific scale factor (hereafter, the “local elongation rate”), *ζ*_*i,j*_. In this version of the model, we ignore promoter-proximal pausing and termination and focus on the gene body, where the elongation signal is easiest to interpret. With these changes, the steady-state density for polymerase occupancy at nucleotide *i* along the body of gene *j* is given by:

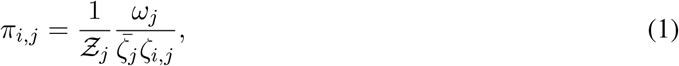

where *ω*_*j*_ is the gene-specific productive initiation rate and the normalization constant for gene *j* of length *N*_*j*_ is given by 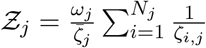.

In turn, the local elongation rate *ζ*_*i,j*_ is defined by a generalized linear function of features along the genome,

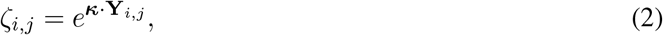

where **Y**_*i,j*_ is the feature vector at site *i* of gene *j* and *κ* is a corresponding vector of real-valued coefficients, whose first element is assumed to be a constant of 1 to accommodate an intercept for the linear function at the corresponding position in *κ*. The use of a single set of coefficients *κ* for all analyzed sites allows sparse information about correlates of elongation rate to be pooled efficiently across many sites and many genes.

As in previous work [24, 25], we assume that the NRS read counts *X*_*i,j*_ for nucleotide *i* of gene *j* are generated by a Poisson process, conditional on the steady-state density *π*_*i,j*_. In particular, we assume that 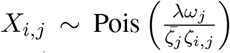, where *λ* is a scale parameter for sequencing read depth. Thus, the expected NRS read counts in gene *j* are proportional to the read depth and the productive initiation rate *ω*_*j*_ and inversely proportional to the gene-wide and local elongation rates 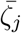 and *ζ*_*i,j*_. The parameters *λ, ω*_*j*_, and 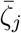, however, are nonidentifiable from steady-state data; only the compound parameter 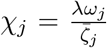 can be estimated from the data (see [24] for discussion). Thus, 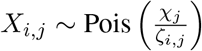.

With these assumptions, the joint log likelihood function for *M* independent genes is given by:

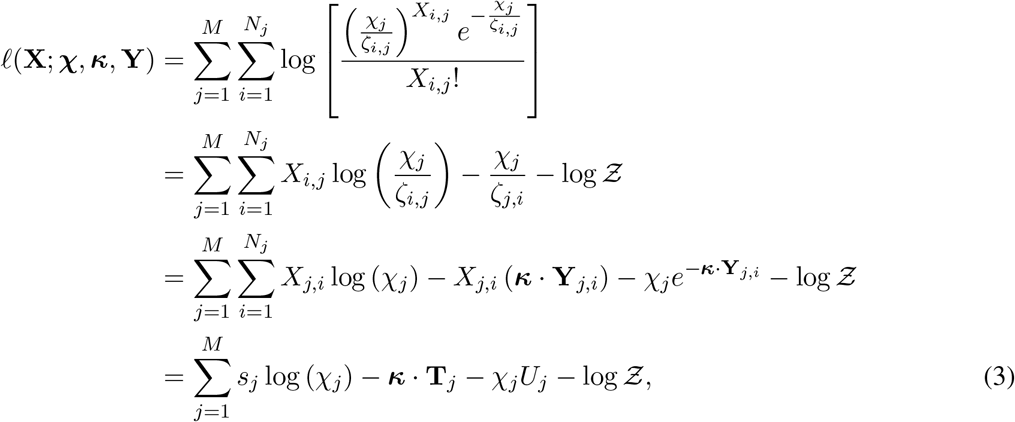

where 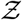 does not depend on the free parameters and can be omitted in optimization, *s*_*j*_ is the sum of read counts for gene 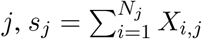, and **T**_*j*_ and *U*_*j*_ are defined analogously as,

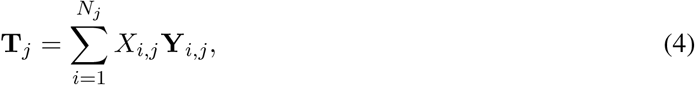

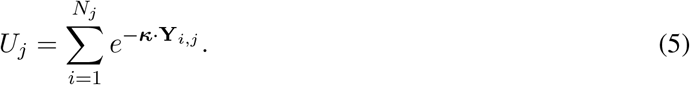

This joint log likelihood can be maximized easily by gradient ascent, in a manner similar to standard Poisson regression. The partial derivative with respect to the *n*th component of *κ* is given by,

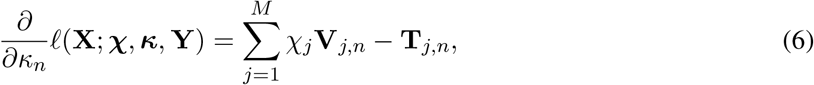

where the final subscript *n* indicates the *n*th element of a vector, and **V**_*j*_ is defined as,

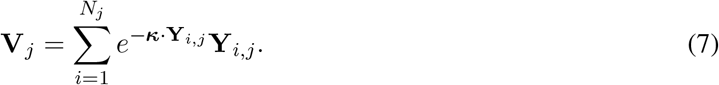

For a given value of *κ*, the maximum for *χ*_*j*_ can be determined analytically as,

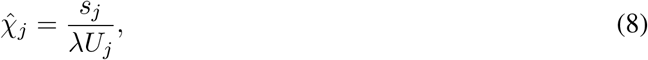

where *λ* is approximated as the average read depth across all genes (see [24]). Thus, the gradient ascent algorithm iteratively improves estimates of *κ* and, on each iteration, fully optimizes *χ*_*j*_ conditional on the other parameters. Notice that the sufficient statistics *s*_*j*_ and **T**_*j*_ need only be computed once, in pre-processing, but *U*_*j*_ and **V**_*j*_ must be recomputed on each iteration of the gradient ascent algorithm. We used a learning rate of 10^*−*7^ (i.e., the multiplier for the gradient on each iteration) for gradient ascent.

### Penalized likelihood extension

In the case of a high-dimensional feature vector (i.e., with the *k*-mer or combined versions of the model) we augmented the log likelihood with a sparsity penalty. We experimented with L1 (lasso), L2 (ridge regression), and combined L1/L2 (elastic net) penalties but found the L1 version to work best in this setting. In this version, the objective function to maximize is (cf., equation 3),

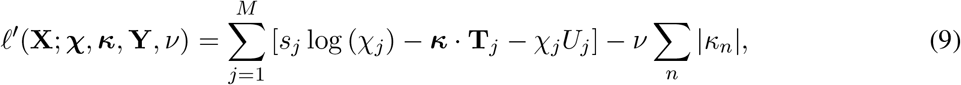

where *ν* is a hyperparameter determining the strength of the penalty and the final sum is over all features. Here, the partial derivative with respect to the *n*th component of *κ* is given by,

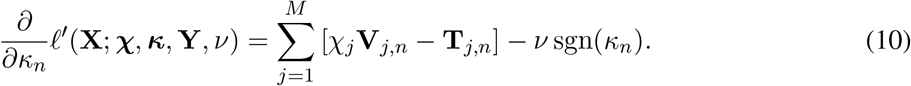

We determine a value for the hyperparameter *ν* separately for each analysis by cross validation. Specifically, we set *ν* to the value that maximized the Akaike Information Criterion (AIC) for held-out testing data, using 80% of the data for training and 20% for testing (see **Supplementary Fig. S7**).

### Extension allowing for sequence bias

We also implemented a version of the *k*-mer model that allows for some, potentially unknown, source of nucleotide bias at the 3^*′*^ ends of aligned PRO-seq reads, and attempts to find *k*-mer associations relative to that bias. The idea behind this model is to see if larger *k*-mer associations persist even if the nucleotides at the apparent active site are somehow biased by the the protocol (see **Supplementary Fig. S9C**).

This version of the model allows for arbitrary relative frequencies of 3^*′*^ nucleotides *π*_A_, *π*_C_, *π*_G_, and *π*_T_, which in practice are pre-estimated from the bulk distribution of 3^*′*^ nucleotides in the PRO-seq reads. They are accommodated in the model by replacing the single read-depth scale parameter *λ* with a separate scale factor for each nucleotide, *λ*_A_, *λ*_C_, *λ*_G_, and *λ*_T_, such that for each base *b* ∈ {A, C, G, T}, *λ*_*b*_ = 4*λπ*_*b*_. For mathematical convenience, we then reparameterize using 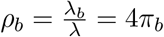.

After this generalization, the (unpenalized) log likelihood becomes (cf. equation 3),

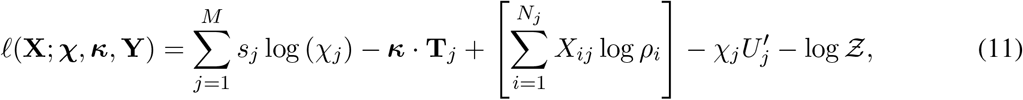

where we use the notation *ρ*_*i*_ to indicate the value of *ρ*_*b*_ corresponding to the nucleotide at position *i*, and where 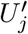 is like *U*_*j*_ (cf. equation 5) but has its terms weighted by the corresponding *ρ*_*b*_ parameters:

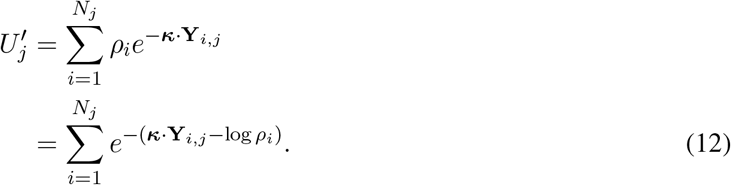

The effect of this model is to add to each dot product *κ* · **Y**_*i,j*_ a quantity of − log *ρ*_*i*_. Thus, nucleotides *b* that are over-represented (with *ρ*_*b*_ *>* 1) are penalized, whereas nucleotides that are under-represented (with *ρ*_*b*_ *<* 1) are rewarded. As a result, *k*-mer associations that simply reflect the background distribution are down-weighted and ones that represent departures from that distribution are up-weighted. If the nucleotides are uniformly distributed (with *ρ*_*b*_ = 1), the model collapses to the original version.

### Smoothing filters for genomic features

As specified, the model requires that any influence on the elongation rate at nucleotide *i* of gene *j* must be captured by the feature vector for the same position, **Y**_*i,j*_. Some features, however, appear to have broader effects that spread out to adjoining nucleotide positions. For example, 3^*′*^ and 5^*′*^ splice sites are narrowly defined at a few nucleotide positions, but metaplots of NRS data suggest that their effects on elongation rate extend for as much as a hundred nucleotides (see, e.g., **Supplementary Fig. S6A–C**), likely because physical interactions between Pol II and the spliceosome can occur over a fairly broad region.

To address this problem, we introduced a preprocessing device called a *smoothing filter* that can be applied to any genomic feature to cause its influence to be distributed to adjacent nucleotide positions. Even for features that are not narrowly defined at a few nucleotides, smoothing filters can be useful in compensating for different levels of genomic resolution across features, for example, to put different ChIP-seq data sets on the same genomic scale.

Formally, a filter *F*_*r,σ,δ*_ is a function defined by three parameters: a *radius* of application *r*, a *smoothing bandwidth σ*, and an *offset δ*. Applying a filter requires replacing each (scalar) covariate 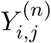 with a filtered version, 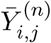 such that,

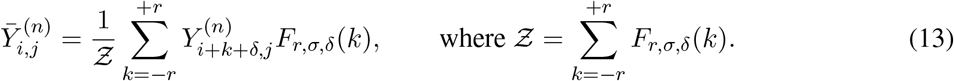

The filter *F*_*r,σ,δ*_(*k*) can take a variety of functional forms, and simple filters can be composed to create more complex ones (see [24]). In this work, however, we found it most useful to work with a Gaussian filter,

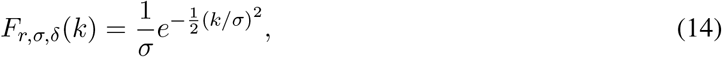

and a generalized filter,

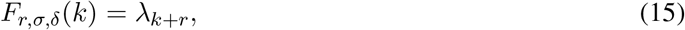

which is defined by a vector of nonnegative scale factors, ***λ*** = {*λ*_0_, …, *λ*_2*r*_} that were estimated from metaplots of PRO-seq data centered on the feature of interest. We used the generalized filter for the 3^*/*^ and 5^*′*^ splice-site features, and the Gaussian filter for the ChIP-seq-based features, including the histone modifications and CTCF (*r* = 400 bp, *σ* = 100 bp; see examples in **Supplementary Fig. S6**). We also applied a Gaussian filter (with *r* = 500 bp, *σ* = 200 bp) to the stem-loop feature, matching it to the corresponding metaplot.

### Standardization of feature values

As with any linear modeling application, the epigenomic and *k*-mer features needed to be standardized to allow the estimated coefficients to be on the same scale and comparable with one another. We used the simple approach of shifting and rescaling the values for each feature to have a mean of zero and standard deviation of one. Some features were defined for only a subset of genomic positions (such as DNA methylation, which was relevant only at CpG dinucleotides); in these cases, the defined values were first standardized and the remaining positions were subsequently assigned values of zero, to ensure that they had no positive or negative effect in the linear model.

Standardization of the indicator features for *k*-mers led to a computational problem that required special attention. Prior to standardization, these features had values of zero at the vast majority of genomic positions, but after standardization these zeroes were converted to negative real values. As a result, the calculation of the *U*_*j*_ and **V**_*j*_ values needed for each iteration of the gradient ascent algorithm (equations 5 and 7) became considerably more laborious. We addressed this problem by first calculating the *U*_*j*_ and **V**_*j*_ values from the unstandardized feature values (containing mostly zeroes) and using a linear transformation to convert them to the corresponding values for the standardized features. In this way, the speed of processing the unstandardized values could be maintained while properly considering the effects of standardization.

### SimPol simulator

The SimPol (“Simulator of Polymerases”) program tracks the independent movement of RNA polymerases along the DNA templates of a large number of cells (**Fig. 2A**). As detailed in ref. [25], the original program accepts user-specified parameters for the initiation pause-escape, and elongation rates, as well as the number of cells being simulated, the gene length, and the total time of transcription. For this study, we modified the simulator to accept a vector of position-specific elongation rates, which could be pre-computed based on covariates (see below). We also omitted the pause-escape portion of the simulation model. Based on the specified parameters, the simulator simply allows each polymerase to move forward or not at the specified rates, in time slices of 10^*−*4^ minutes, assuming at most one movement per time slice, and prohibiting movement if another polymerase blocks forward progress. The program was run until polymerase occupancy along the gene body reached equilibrium (20 min simulated time), and then the empirical density was output as a file in csv format. From this output, an accompanying R script was used to sample synthetic NRS read counts at each nucleotide position (as in [25]), with a target mean read depth equal to that observed in the real PRO-seq data for K562 cells (**Supplementary Fig. S1B**).

### Generation of synthetic NRS data

For the epigenomic simulations, we used SimPol to simulate 10 replicates of 100 TUs of length 10 kbp in 5,000 cells. In each replicate, gene-specific initiation rates were sampled from real data for K562 cells [25], by rescaling the estimated *χ* values as initiation rates *α* having a median of 1 event per min. We selected six representative epigenomic features from real data for K562 cells: CTCF binding sites, four histone marks, and RNA stem-loops (see **Fig. 2B**). To generate synthetic covariates, we sampled combinations of these covariates from the real data in 1 kbp blocks (**Supplementary Fig. S1C & D**), ensuring that their correlation structure was preserved. We then generated feature-specific local elongation rates according to the same generalized linear model used for inference, but with the addition of Gaussian noise. Specifically, we assigned a *κ* coefficient to each feature and set it equal to a value estimated in a preliminary analysis of the K562 data, and then we set the local elongation rate for each position *i* to be *ζ*_*i*_ = exp(*κ* ·**Y**_*i*_)+*δ*_*i*_, where *δ*_*i*_ ∼ 𝒩 (0, 0.1). The vector of simulated *ζ*_*i*_ values was then passed to SimPol for simulation of polymerase movement and NRS read counts (above). We also experimented with a version of these simulations without Gaussian noise (**Supplementary Fig. S2A–D**).

The simulations for the 5-mer model were similar, but in this case we simulated 200 TUs in each replicate, owing to the high-dimensional feature vector. We randomly sampled 100 5-mers and assigned them coefficients ranging from −0.3 to 0.3, while the remaining 5-mers were assigned a coefficient of 0. The initiation rates and position-specific elongation rates were determined as above, but without the addition of Gaussian noise.

### Analysis of real data

We acquired PRO-seq data sets for K562, CD14+, Hela-S3, and MCF-7 cell lines from published sources [26, 31, 40, 41] and processed them using the proseq2.0 pipeline (https://github.com/Danko-Lab/proseq2.0) [49]. The data were processed exactly as described in ref. [25]. Briefly, mapping was performed with human genome assembly GRCh38.p13. The 3^*′*^ ends of mapped reads—which we take to represent the active sites of transcriptionally engaged polymerases—were recorded in bigWig files and used for analysis. Gene annotations were downloaded from Ensembl (release 99) in GTF [50]. Annotations of protein-coding genes from the autosomes and sex chromosomes were used, excluding overlapping genes on the same strand. DENR [31] was applied separately to each data set to select dominant pre-mRNA isoforms and estimate corresponding expression levels. Genes with a DENR-estimated abundance of *<*10 TPM (Transcripts Per Kilobase Million) were removed.

We took several measures to refine gene annotations, select regions for analysis, and remove signals not representative of elongation rates. First, we refined the annotated TSS positions using cell-type-matched NRS data. For the K562 data, we used available CoPRO-cap (Coordinated Precision Run-On and sequencing with 5^*′*^ Capped RNA) data [30] to re-position the TSS within a region −1500 bp to +1500 bp of the annotated TSS selected by DENR. In the other three cell lines, we performed a similar refinement using the 5^*′*^ ends of aligned PRO-seq reads (as in [25]). Second, we conservatively defined the “gene body” for each gene as the interval from 2250 bp downstream of the refined TSS (a distance we verified was sufficient to eliminate all pause peaks) to 250 bp upstream of the annotated TTS. Gene bodies *<*6 kbp were omitted. Finally, to eliminate potential internal TSSs—which create PRO-seq peaks not representative of gene-body elongation rates—we used GRO-cap (Global Nuclear Run-On and sequencing with 5^*′*^ Capped RNA) [33] or PRO-cap (Precision Run-On and sequencing with 5^*′*^ Capped RNA) [51] data where available, as well as dREG [32] predictions from the primary PRO-seq data. Specifically, we masked out predicted dREG peaks and 2-kbp intervals centered on GRO-cap or PRO-cap peaks (with read counts *>*10) (see example in **Supplementary Fig. S4**). In later analyses (with MCF-7), we omitted the less reliable dREG filter and used PRO-cap only. The final data sets consisted of 6,391 genes for K562, 5,336 for CD14+, 6,657 for HeLa, 6,193 for MCF-7. For the comparison of cell lines, we considered the intersection of these sets, which consisted of 3,716 genes.

To eliminate the shared “U-shape” pattern along gene bodies, we first merged the data for all genes after standardizing their lengths to a common scale and normalizing the raw read counts by dividing them by the median for each gene (see **Supplementary Fig. S5**). We then smoothed the merged data using the LOESS method, creating a gently U-shaped curve with an average height of one. We then adjusted the raw read counts by dividing by the height of the LOESS curve at the corresponding positions, ensuring that the data were flat on average. These adjusted read counts served as the inputs *X*_*i,j*_ for our model.

For the analysis of pausing locations within gene bodies (**Fig. 3E** and **Supplementary Fig. S9A**), we partitioned each gene body into 200 bp intervals, summed the PRO-seq read counts within each window, and identified the top 5 intervals for each gene as putative pausing locations.

### Test for ligation bias

To test for a nucleotide bias in ligation, we leveraged the particular adapter design used for our CD14+ PRO-seq library, as follows. These adapter designs, denoted as RA3 and RA5, incorporate a UMI consisting of random bases (NNNNNN) at the 3^*′*^ end of the 5^*′*^ adapter (see **Supplementary Fig. S10C**). In this configuration, the 3^*′*^ end of a UMI mirrors the 3^*′*^ end of an insert, allowing for a natural comparison of the nucleotide composition of insert-adapter dimers with no-insert adapter dimers. We first established that the 3^*′*^ ends of the synthesized UMIs have a uniform distribution of nucleotides (**Supplementary Fig. S10D**). Next, we examined the 3^*′*^ ends of UMIs that were either ligated to RA3 (*no inserts*) or inserts (*with inserts*). In a pool of ∼20 million sequenced reads, about ∼90% were insert-adaptor dimers, and ∼3% were no-insert adapter dimers. We found no major difference between the 3^*′*^-most nucleotides in these two sets (**Supplementary Fig. S10E**), suggesting no ligation bias. In particular, the cytosine frequencies in the two sets were nearly identical.

### Enrichment logos for 5-mer candidates

To determine the enrichment logos of the 5-mer model, we clustered them using a *K*-means algorithm. The 5-mers were divided into two groups based on their positive or negative *κ* values, with the top 50 candidates selected for each group. For each group, a *K*-means method with *K* = 2 was applied to cluster 5-mers with similar *κ* values. The enrichment of the sequence logos for each cluster was visualized using *ggseqlogo* [52].

## Funding

This research was supported by US National Institutes of Health grants R35-GM127070 (to AS) and R01-HG010346 (to Charles Danko and AS), and by the Simons Center for Quantitative Biology at Cold Spring Harbor Laboratory. The content is solely the responsibility of the authors and does not necessarily represent the official views of the US National Institutes of Health.

## Conflict of Interest

The authors declare no competing interests.

## Acknowledgments

We thank Charles Danko, Gilad Barshad, David McCandlish, Peter Koo, Shushan Toneyan, Hannah Meyer, and other members of the community at Cold Spring Harbor Laboratory for helpful discussions.

## Notes

### Competing Interest Statement

The authors have declared no competing interest.

