## Supplementary Material for "DNA-sequence and epigenomic determinants of local rates of transcription elongation"

### Supplementary Information for: DNA-sequence and epigenomic determinants of local rates of transcription elongation

### Supplementary Figures

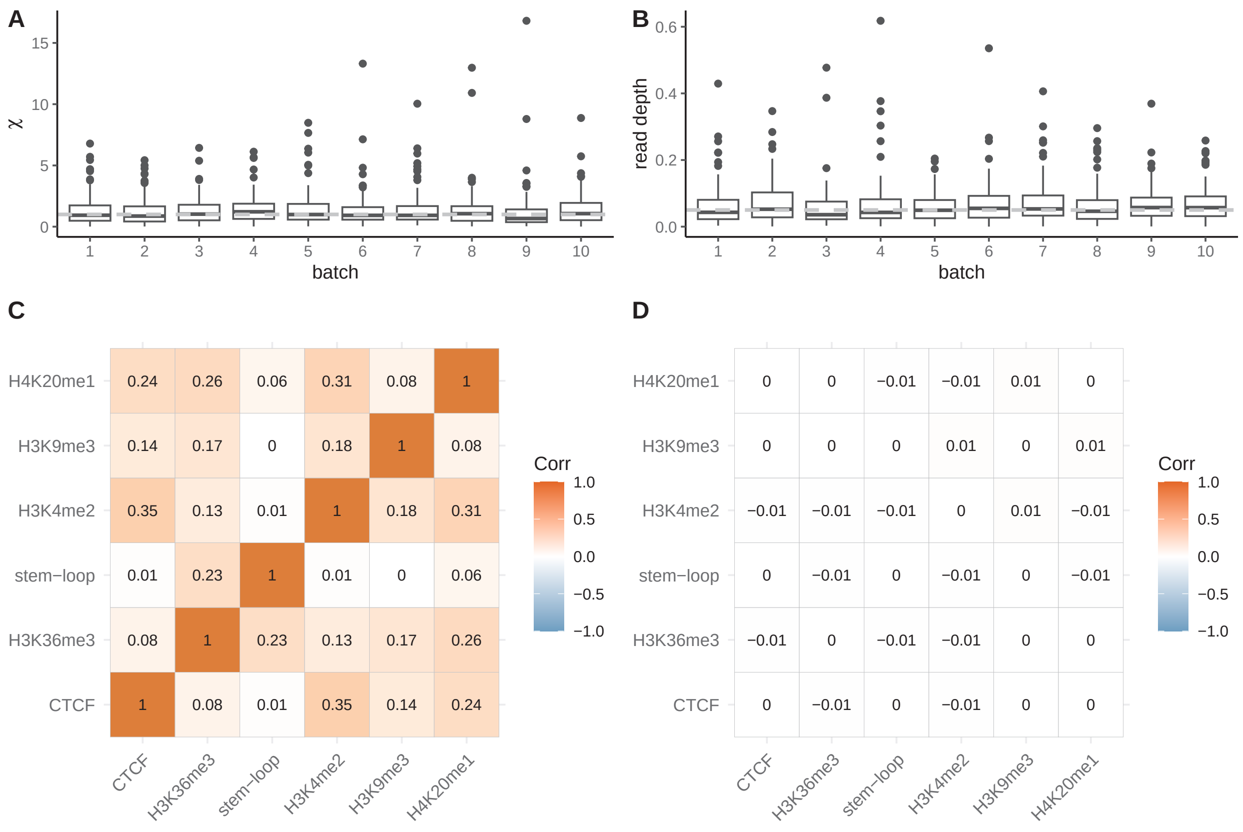


Supplementary Figure S1: **A**. The distribution of productive initiation rates $\chi$ in 10 rounds of simulation sampled from the estimated $\chi$ of real K562 PRO-seq data is shown. The dashed line represents the median of 1 event per minute [1]. **B**. The distribution of read depth in 10 rounds of simulation of synthetic data is displayed. The read depth is decreased to match the read depth of real K562 PRO-seq data [1], with a median value of around 0.05 [2]. **C**. The correlation map of selected features from real K562 data. **D**. The difference in correlation map of selected features between the sampled covariates and the real K562 covariates.

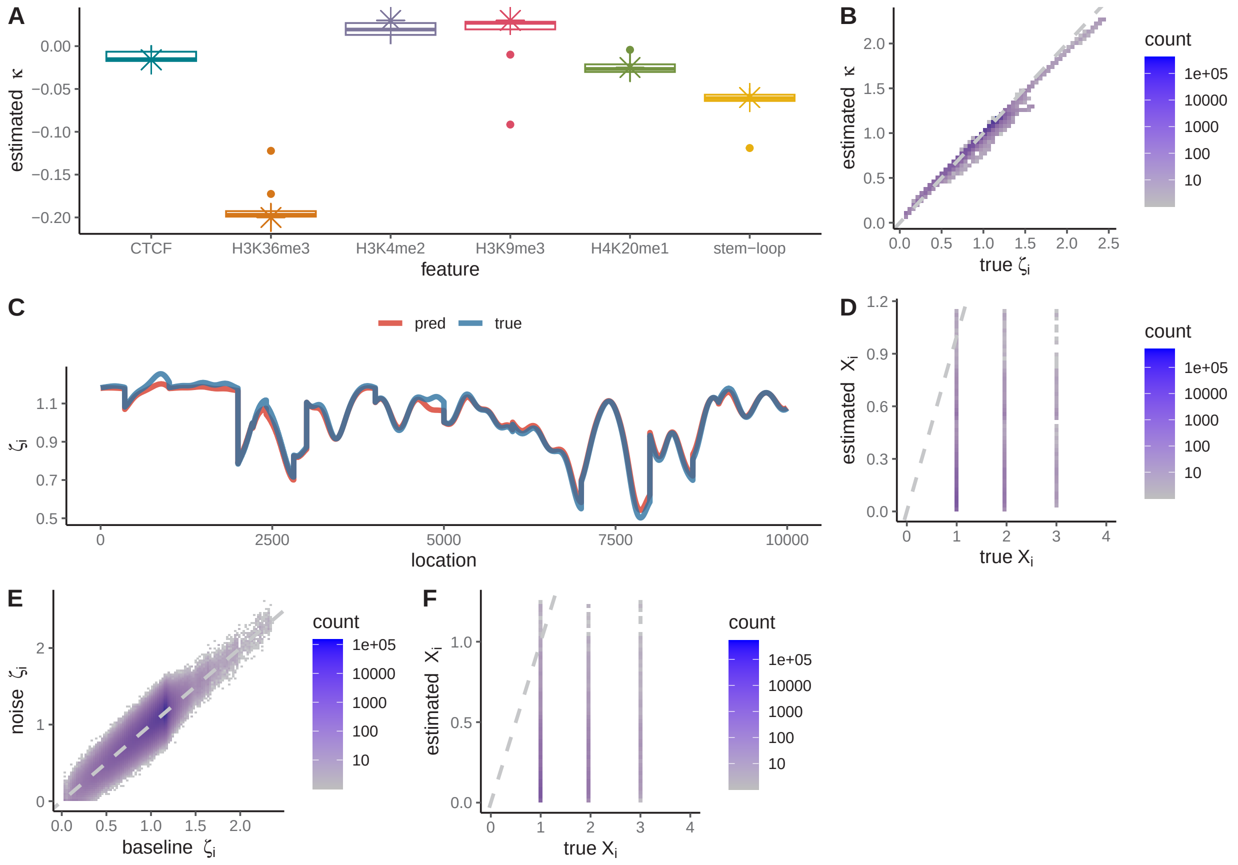


Supplementary Figure S2: **A & B & C & D.** The simulation with an input of baseline $\zeta$ that is exactly determined by a GLM, with no other unmodeled source of variation. **A**. Accuracy of predicted coefficients $\kappa$ using synthetic data over ten rounds of simulation. Crosses indicate the ground truth. **B**. Evaluation of the accuracy of predicted per-nucleotide elongation rates $\zeta_i$ across all TUs by comparing them to the corresponding ground truth, with Pearson's $r^2$ of 0.991 **C**. Evaluation of the accuracy of predicted local elongation rates $\zeta_i$ along an individual TU by comparing them to the corresponding ground truth, with Pearson's $r^2$ of 0.983. **D**. The accuracy of per-nucleotide PRO-seq read count $X_i$ predictions based on selected features is evaluated by comparing them to the synthetic read counts in the held out data of the total 10 rounds of simulation, with Pearson's $r^2$ of 0.26. **E**.The comparison of baseline $\zeta$ that is exactly determined by a GLM and the noise $\zeta$ after introducing Gaussian noise, with Pearson's $r^2$ of 0.749. **F**. The accuracy of per-nucleotide PRO-seq read count $X_i$ predictions in the held out data of the total 10 rounds of simulation with an input of noise $\zeta$, with Pearson's $r^2$ of 0.070.

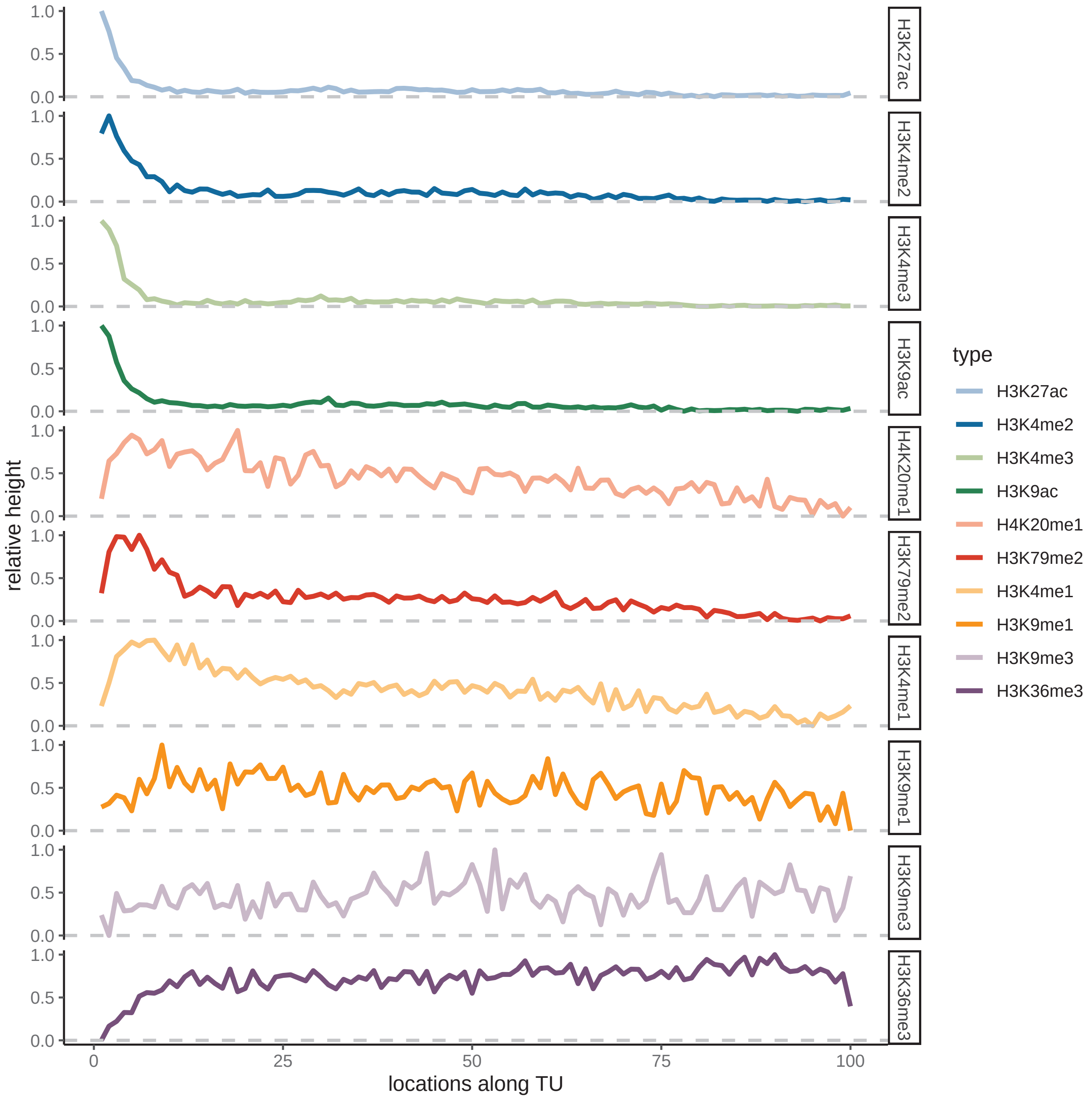


Supplementary Figure S3: **A**. Distribution of multiple histone marks from TSSs to the end of gene bodies. To visualize this distribution, we have scaled the ChIP-seq signals for each histone mark from 0 to 1, which allows us to represent their relative heights on the y-axis.

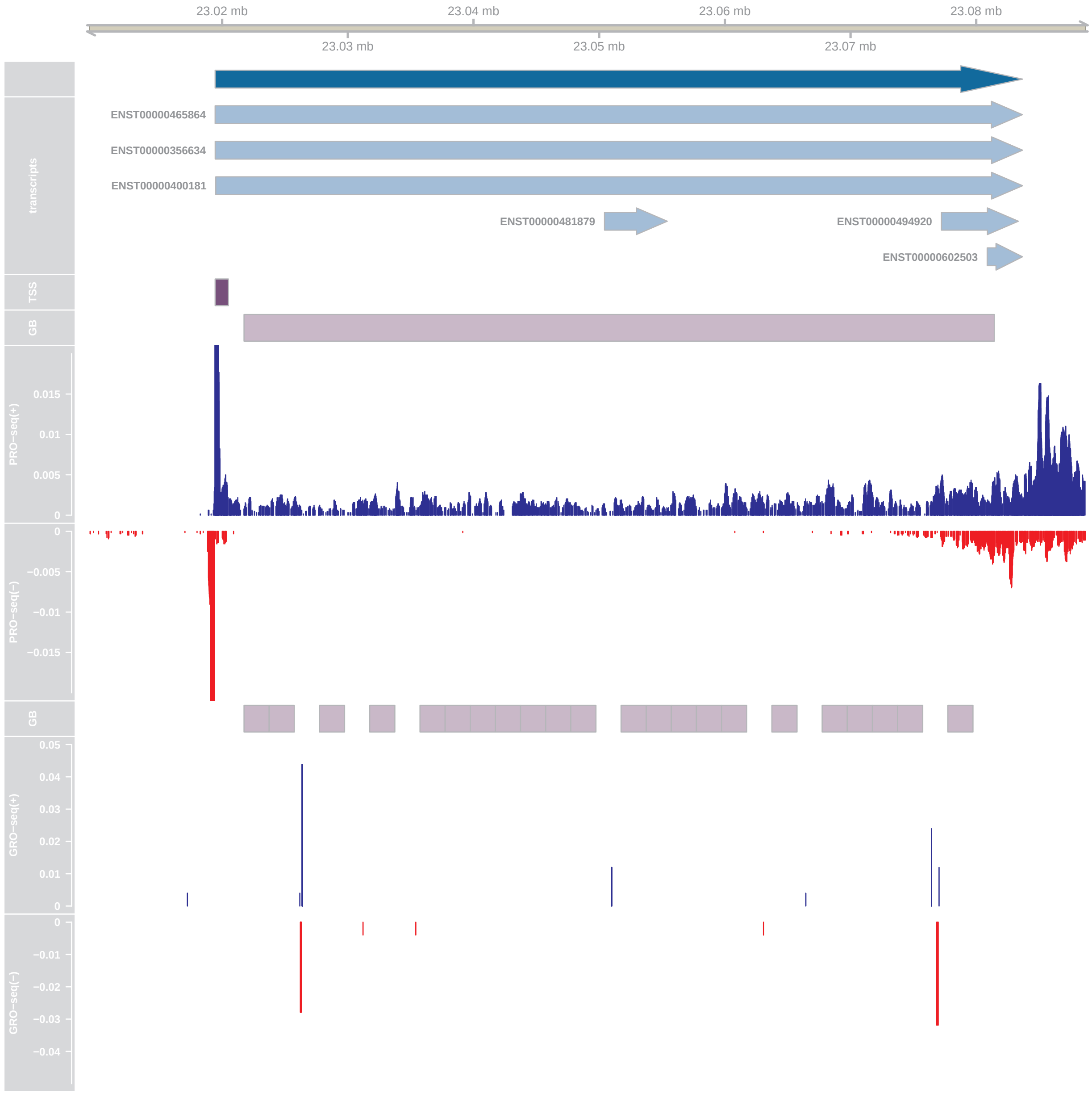


Supplementary Figure S4: The selection of gene body of gene MFSD4B in K562 PRO-seq data.

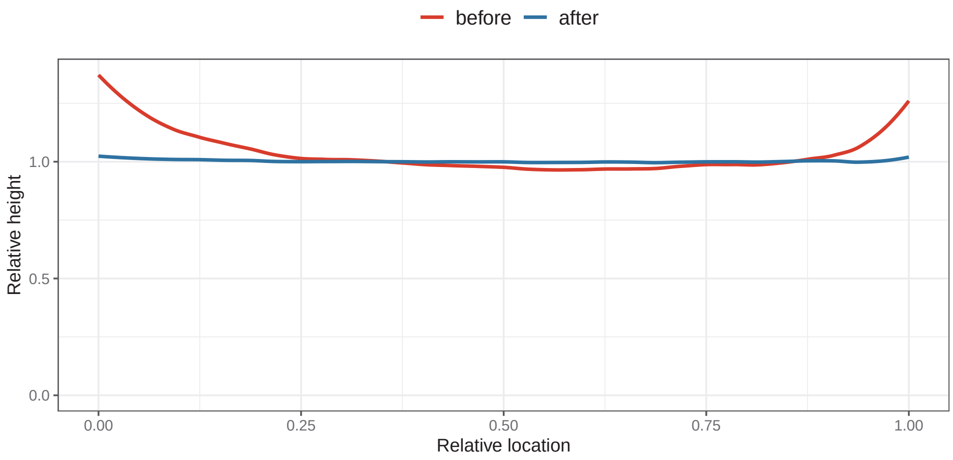


Supplementary Figure S5: A ”U-shape” is evident in the selected gene bodies of K562 PRO-seq data. The red line illustrates the general pattern of the relative height of PRO-seq signals across the relative location of gene bodies for 6,000 genes. The blue line represents the relative height after correcting the ”U-shape.”

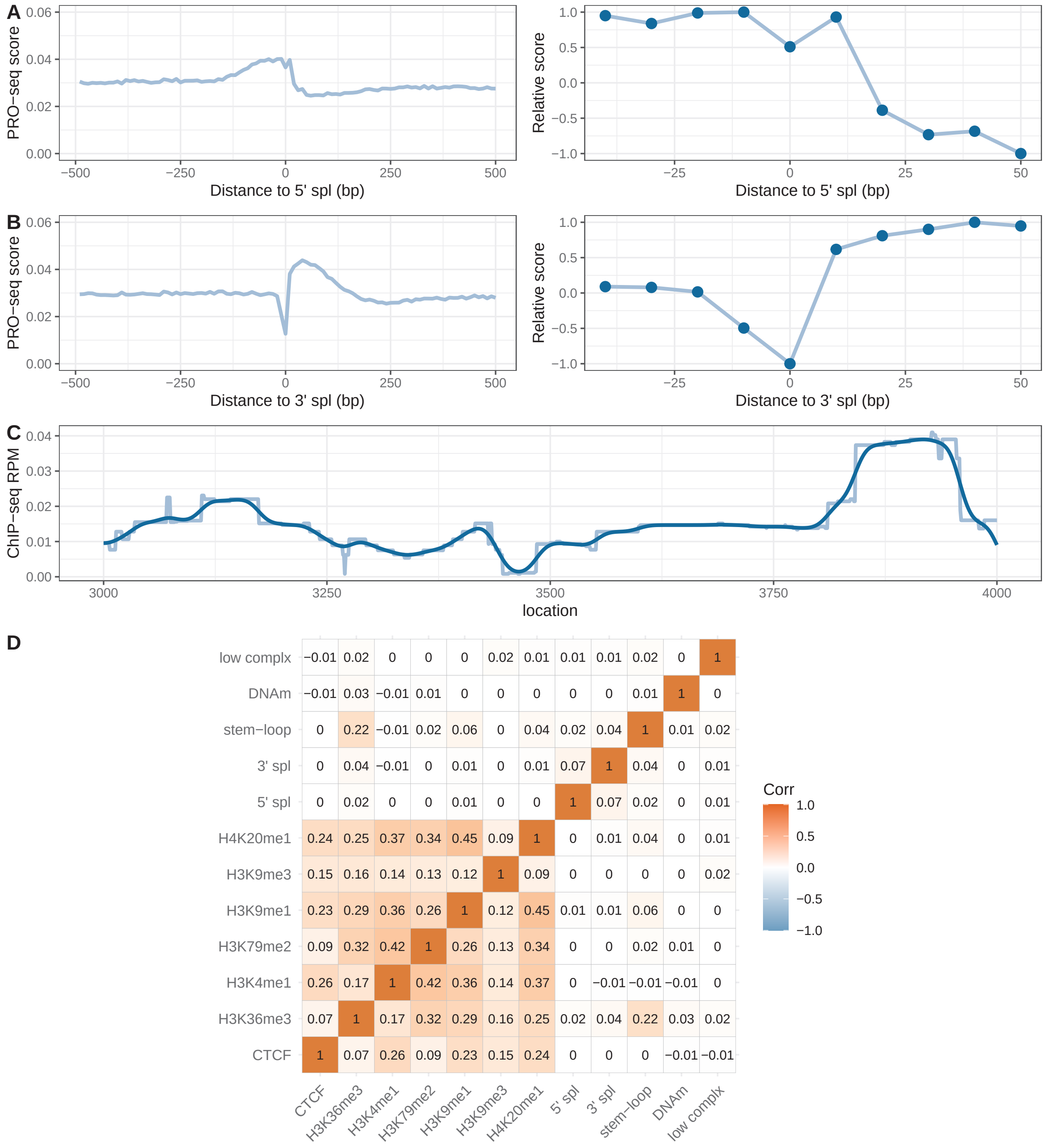


Supplementary Figure S6: **A & B.** The selection of an appropriate filter and its parameters for the 5’ and 3’ splicing sites is accomplished by examining metaplots. The average PRO-seq profile at the 5’ and 3’ splicing sites exhibits discernible effects on the elongation rate within their respective ranges (see *left*). A generalized filter is applied for the $3'$ and $5'$ splice-site features, where the dots indicate the scaling vector (see *right*). **C.** A Gaussian filter is applied to smooth the ChIP-seq signals of histone marks. **D**. The correlation map of all epigenomic and annotated features in the actual analysis of K562 data.

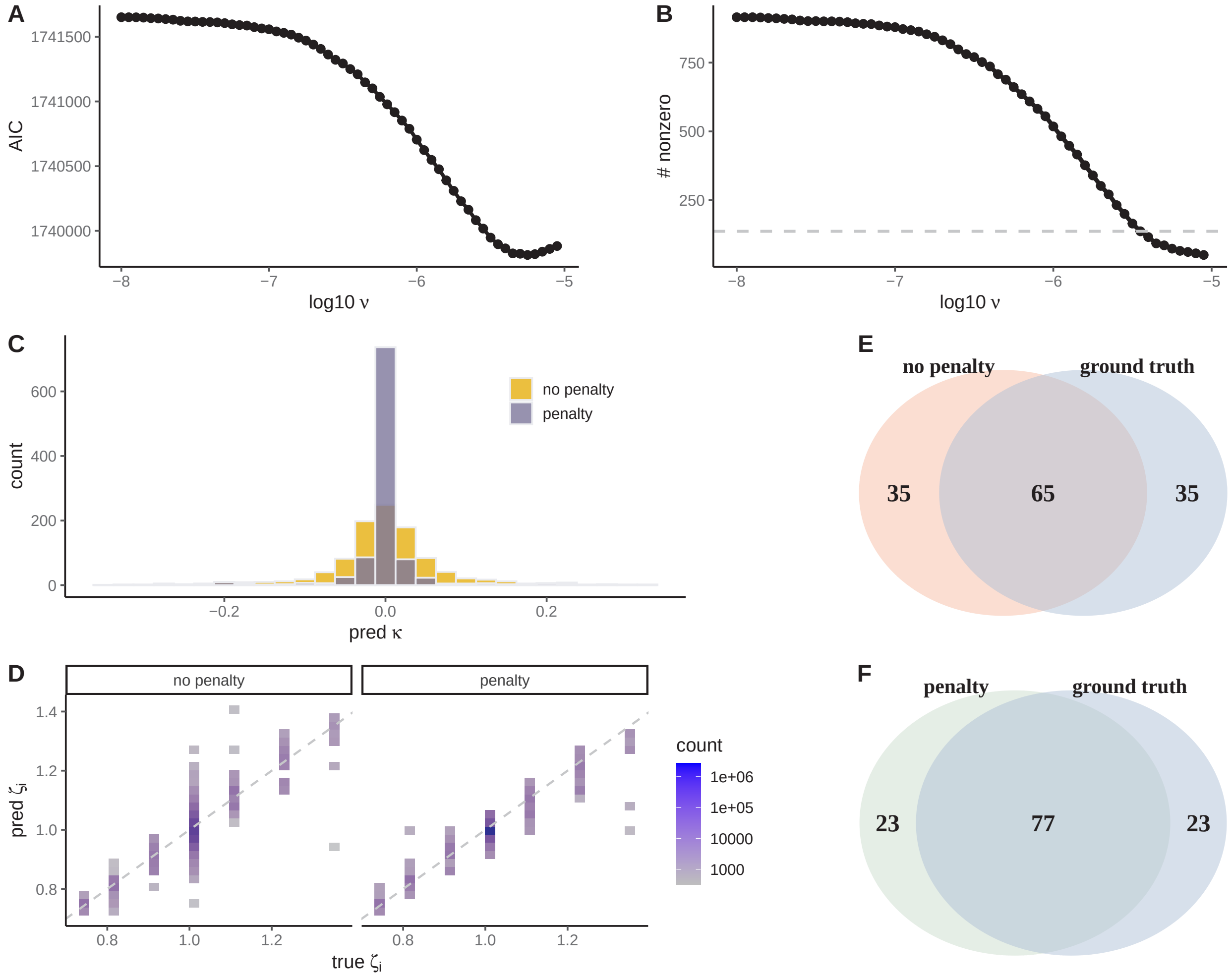


Supplementary Figure S7: Determination of the hyperparameter of L1 regularization in the simulated data. **A**. Evaluation of the predictiveness of each hyperparameter through the computation of Akaike Information Criterion (AIC) values. **B**. The number of non-zero 5-mers by the framework applying L1 regularization. The dashed line represents the number of non-zero 5-mers selected with the hyperparameter corresponding to the lowest AIC value. **C**. Successful shrinkage of parameters after the application of penalty, contrasted with scenarios without penalty. **D**. Accuracy assessment of predicted local elongation rates $\zeta_i$ compared to ground truth, in settings without penalty (left) and with penalty (right), exhibiting Pearson's $r^2$ values of 0.76 and 0.89. **E & F**. Comparison of the top 100 5-mers with the most significant coefficients selected by the model and the ground truth in scenarios without and with L1 penalty, respectively.

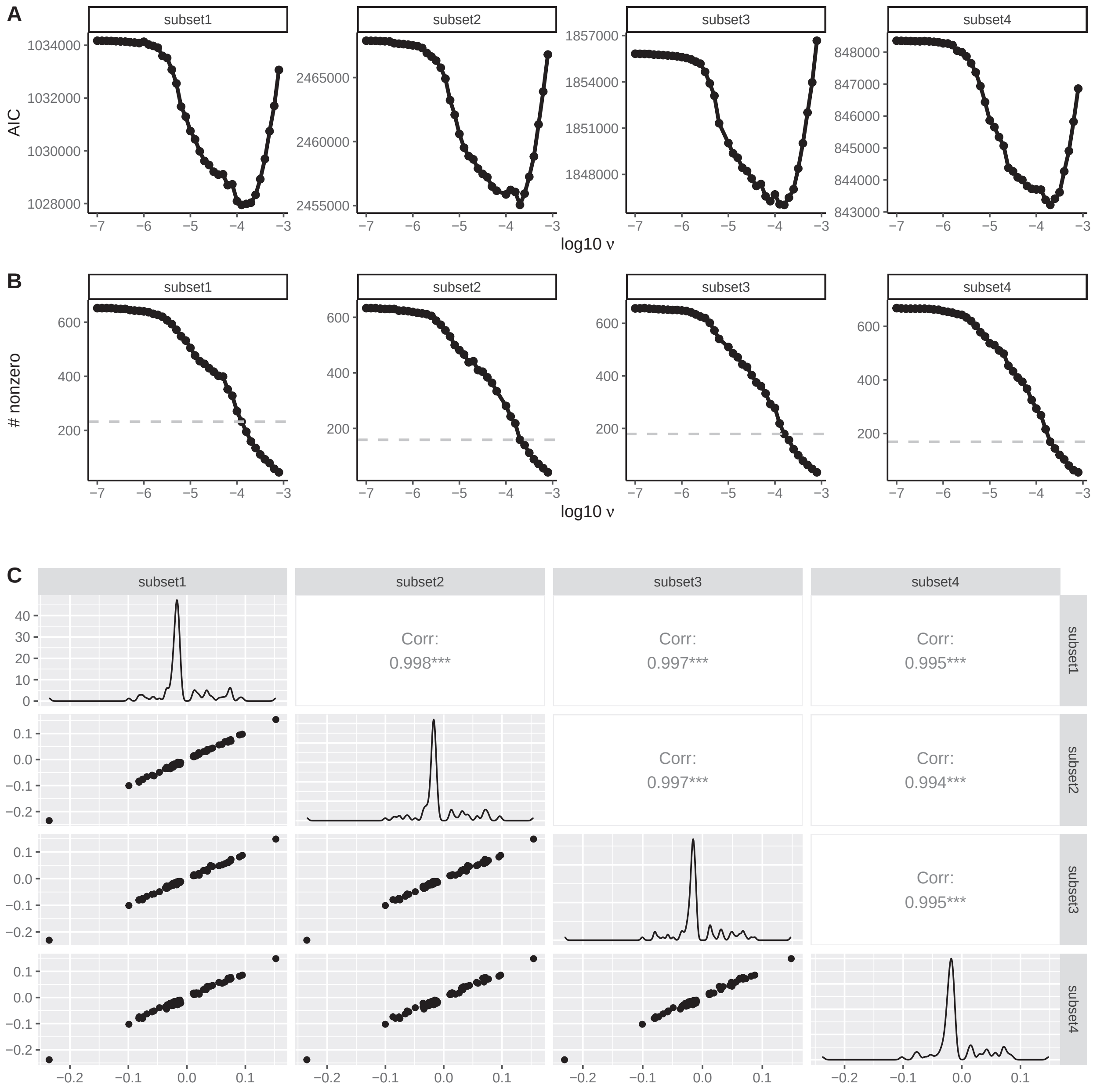


Supplementary Figure S8: **A**. The calculation of AIC values for each hyperparameter across varied data samplings. **B**. The number of non-zero 5-mers by the framework applying L1 regularization across varied data samplings. The dashed line represents the number of non-zero 5-mers selected with the hyperparameter corresponding to the lowest AIC value. **C**. The consistency of the prediction of significant allmers ($N = 105$) across varied data samplings.

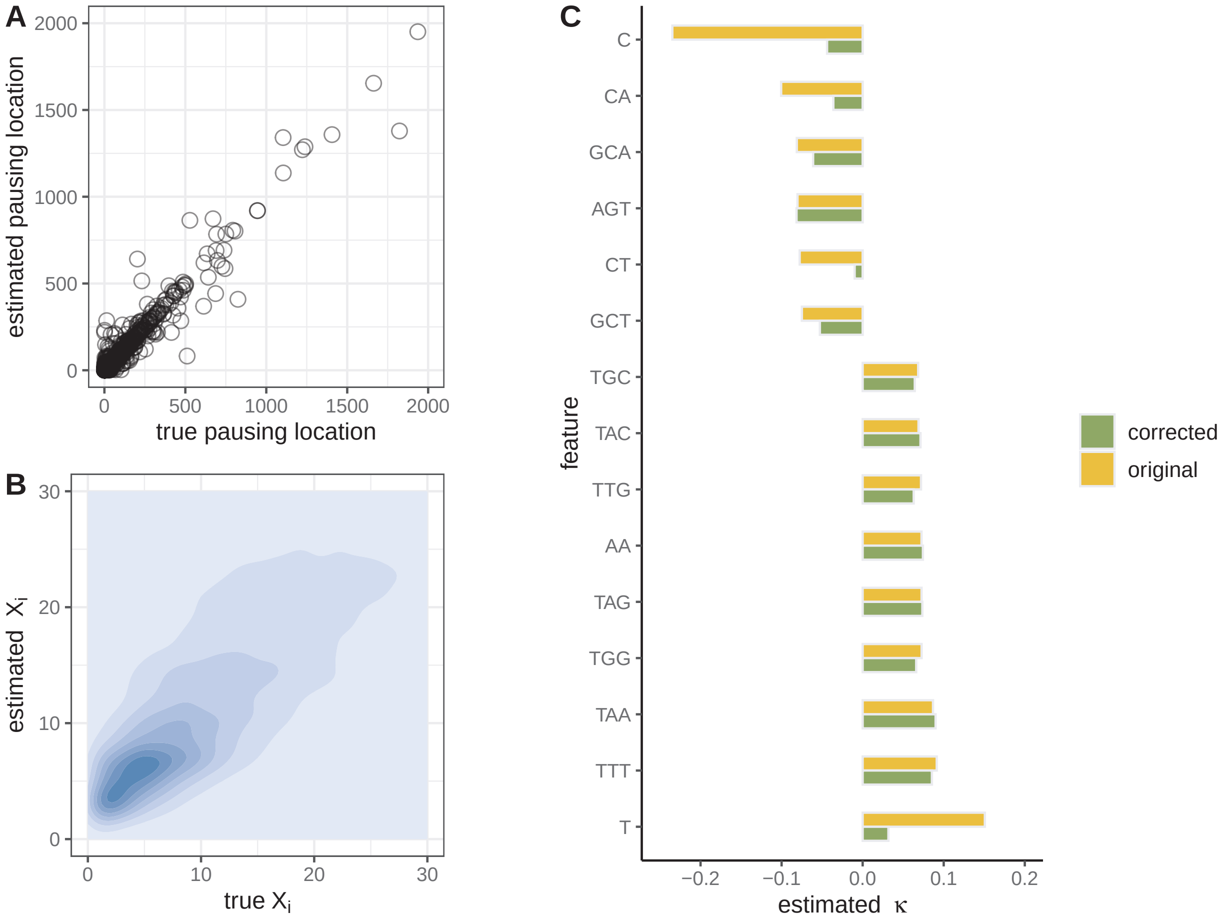


Supplementary Figure S9: The analysis of applying the allmer model to real K562 data. **A**. The model's performance in predicting local pausing sites along the gene bodies of held-out genes was evaluated. The locations of the pausing sites determined by the predicted read counts $X_i$ compared to the true read counts $X_i$, with Pearson's $r^2$ of 0.64. **B**. The accuracy of the predicted read counts $X_i$ was assessed across 1 kbp intervals for all transcribed units (TUs) by comparing them to the corresponding true $X_i$ values, with Pearson's $r^2$ of 0.65. **C**. The comparison of predictiveness among the epigenomic model, $k$-mer model, and the combined model of pausing locations and PRO-seq read counts in 1kbp. Pearson's $r^2$ between estimation and truth is indicated on the y-axis. **D**. Comparative analysis of predicted coefficients $\kappa$ using the original model and a sequence bias model that assumes all genome-wide effects reflect biases.

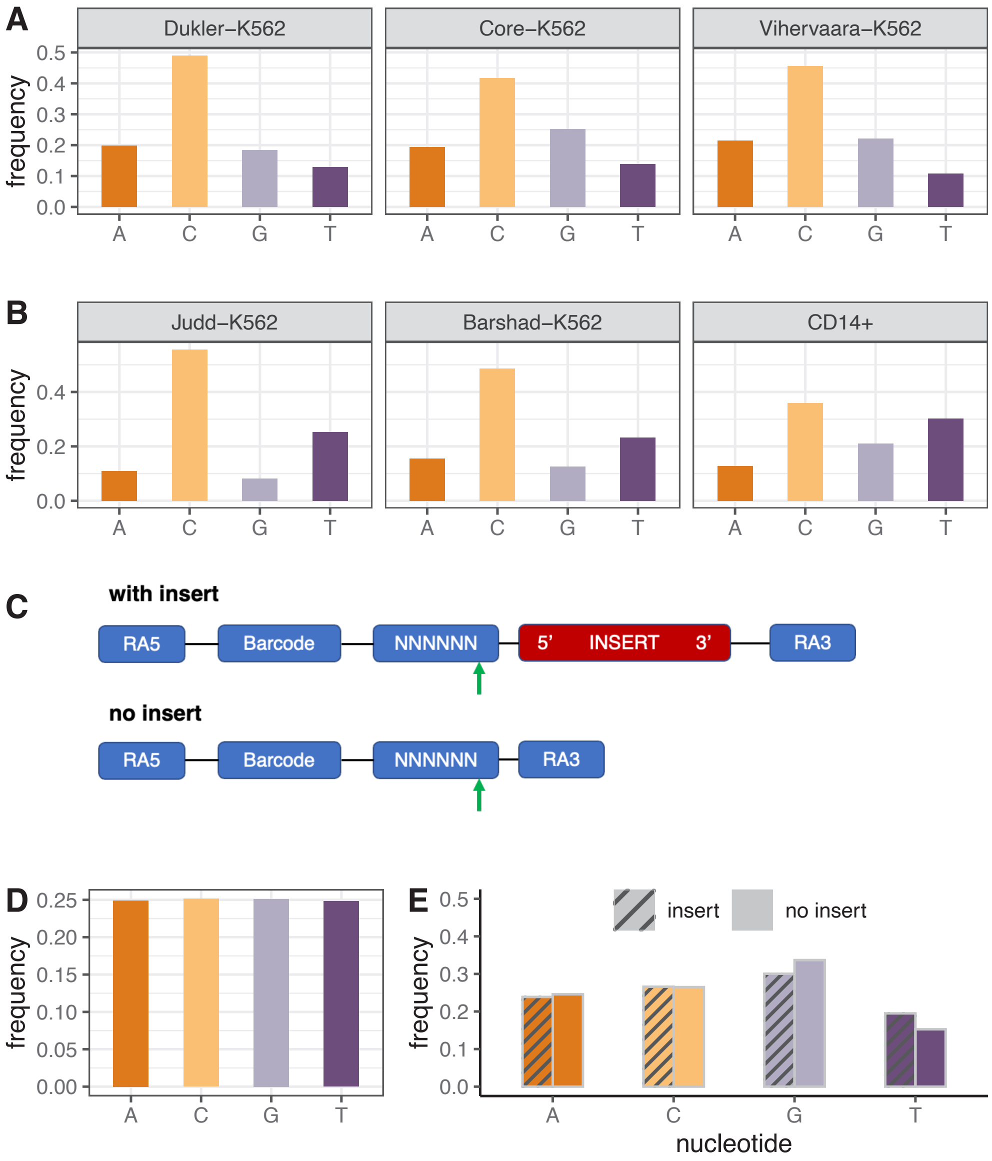


Supplementary Figure S10: **A**. The distribution of the 3′ end base in PRO-seq data from the K562 cell line, acquired through a run-on experiment involving 4 dNTPs with equal concentrations. **B**. The distribution of the 3′ end base in PRO-seq data from the K562 and CD14+ cell lines, acquired through a run-on experiment involving 2 dNTPs. **C**. Examination utilizing CD14+ PRO-seq data with a UMI design ligated to an insert. An illustration depicting the structure of reads, either with or without inserts. The green arrow indicates the 3′ end of UMIs. **D**. The distribution of the 3′ end constitution of UMIs in CD14+ library, with a design to facilitate the examination of potential ligation bias. **E**. Analysis of the distribution of the 3′ end base of UMI, whether with or without successful inserts, indicating an absence of ligation bias towards cytosine.

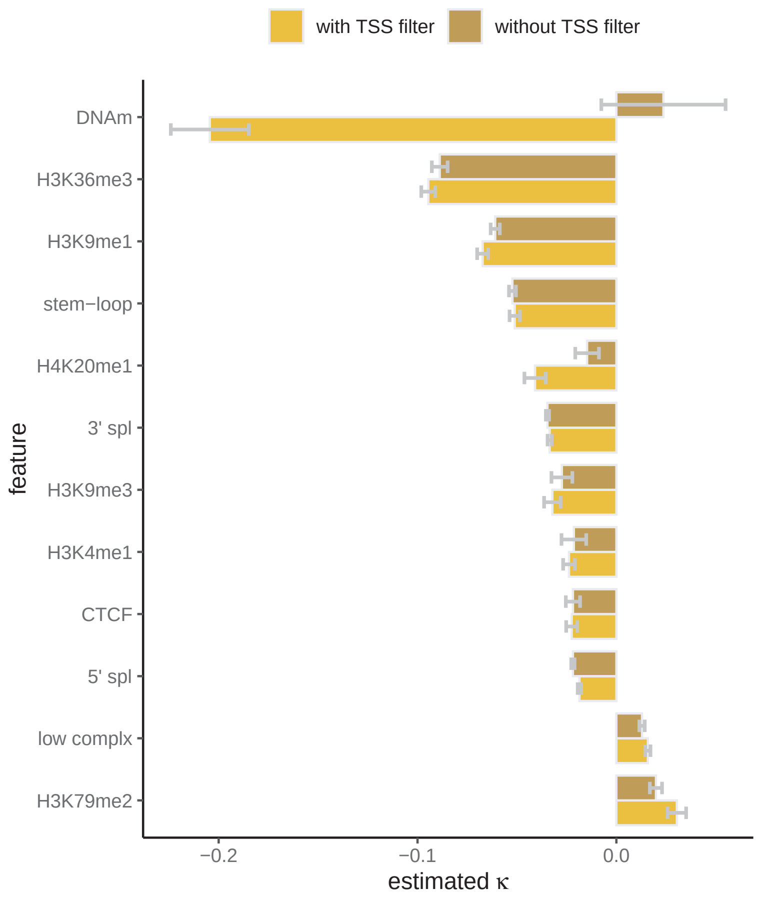


Supplementary Figure S11: Analysis of K562 cells illustrating the impact of potential internal TSSs on the estimation of DNAm coefficients. In the 'without TSS filter' scenario, the entire gene body is included, and in the 'with TSS filter' scenario, internal TSSs based on GRO-seq signals are excluded within the same gene body. Error bars represent the standard deviation of rounds of sampled genes. The most notable difference lies in the DNAm coefficient: 'without TSS filter' shows a positive correlation between DNAm and elongation rate, while 'with TSS filter' shows a negative correlation. This suggests a switch function for DNAm in new initiation within gene bodies, potentially influencing the true correlations between DNAm and elongating RNAP.

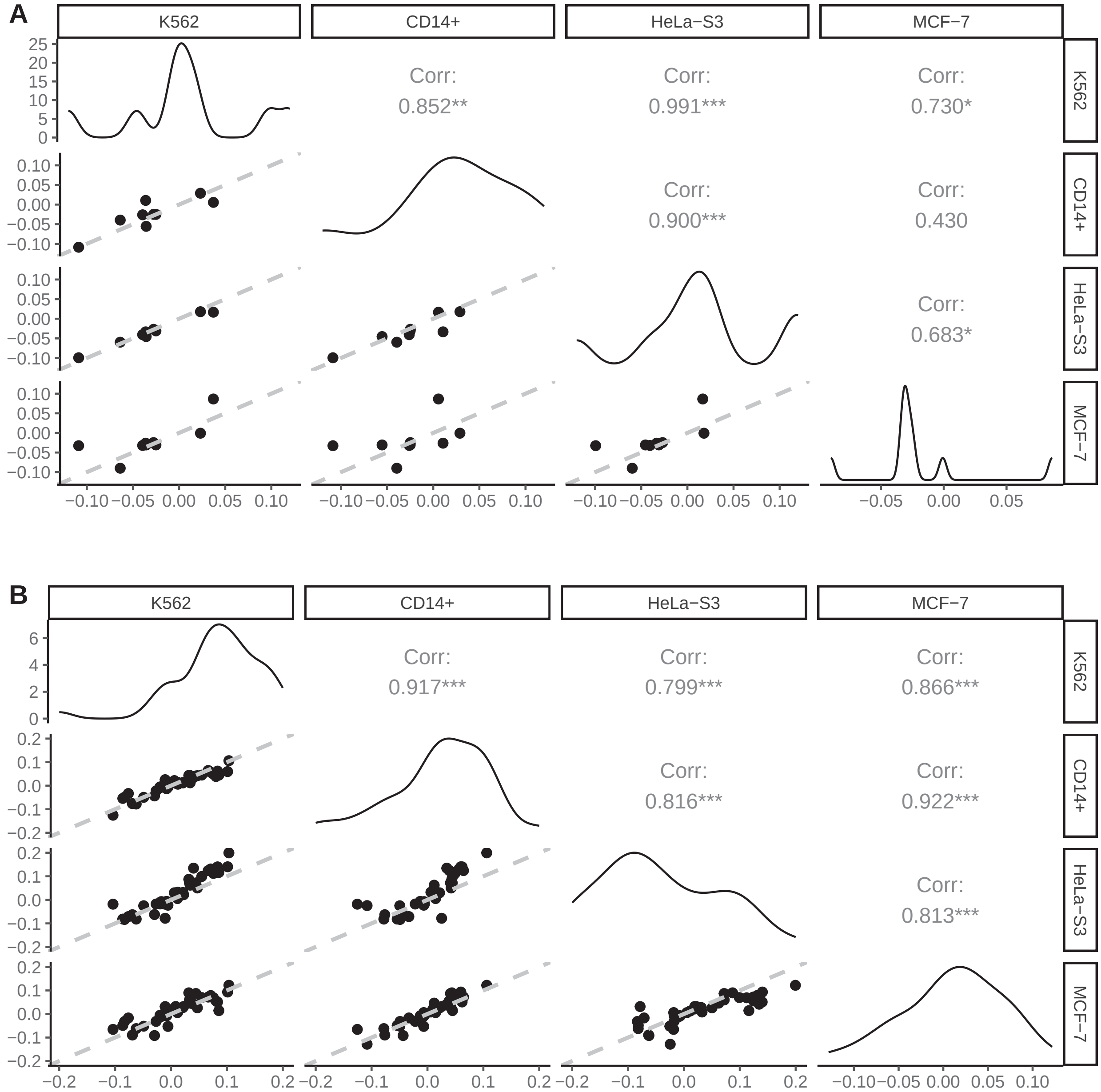


Supplementary Figure S12: **A & B**.The correlation between the estimated $\kappa$ values of epigenomic features and non-zero allmers (N = 45) across four cell lines.

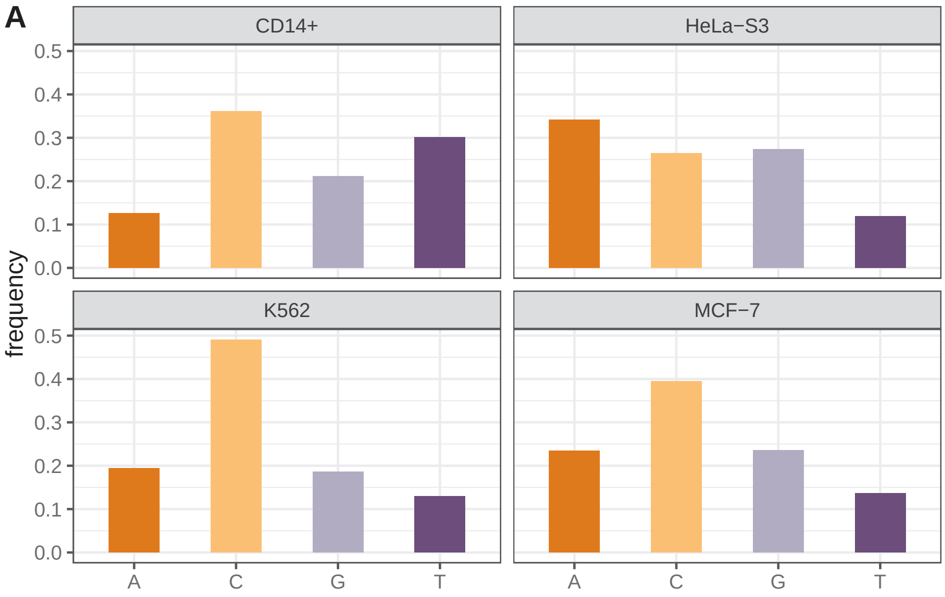


Supplementary Figure S13: **A**. The distribution of the 3′ end base in PRO-seq data across for four cell lines.

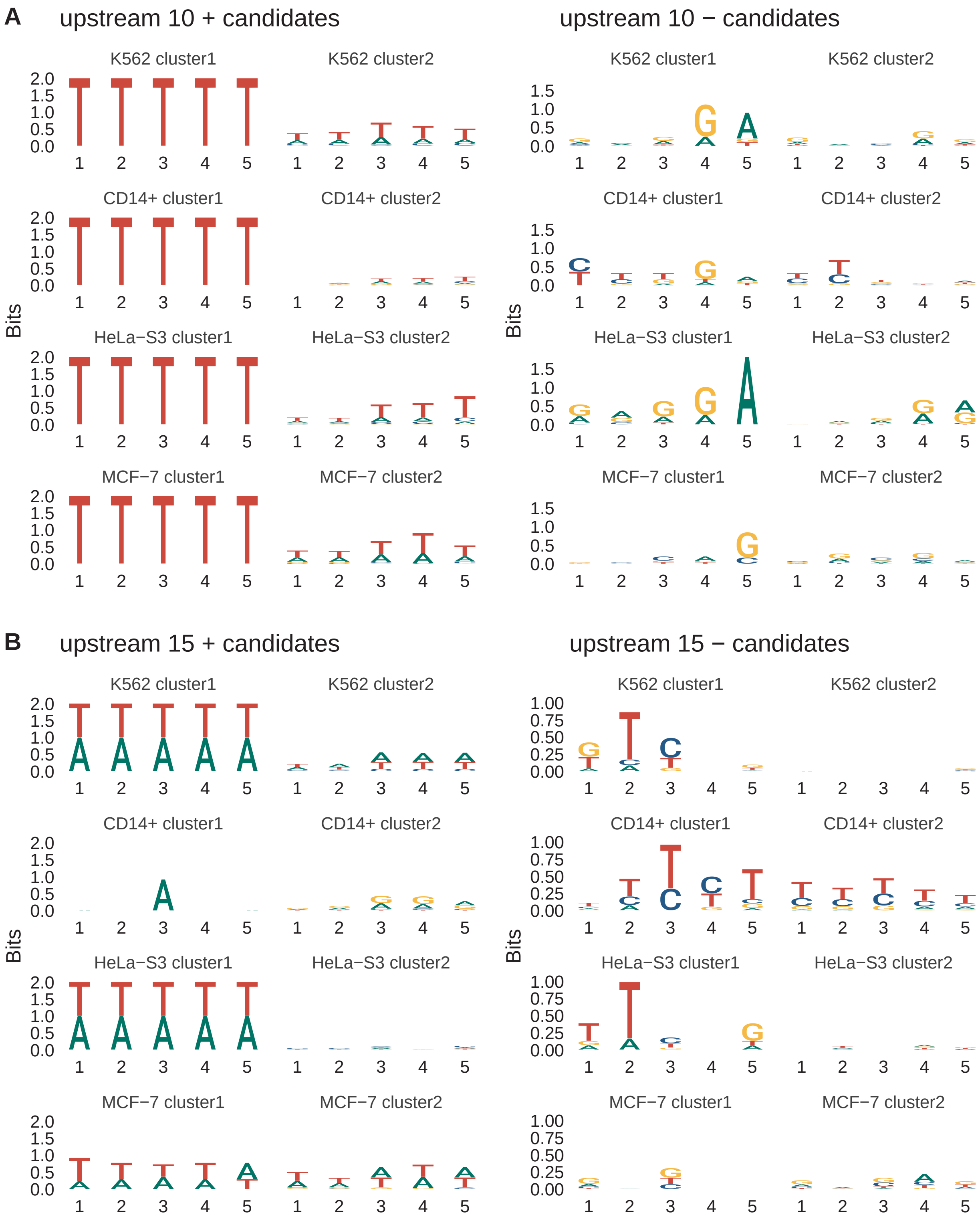


Supplementary Figure S14: **A & B**.Comparison of enrichment logos illustrating the top positive or negative 5-mers in the upstream 10-nucleotide and 15-nucleotide regions along gene bodies for four cell lines.

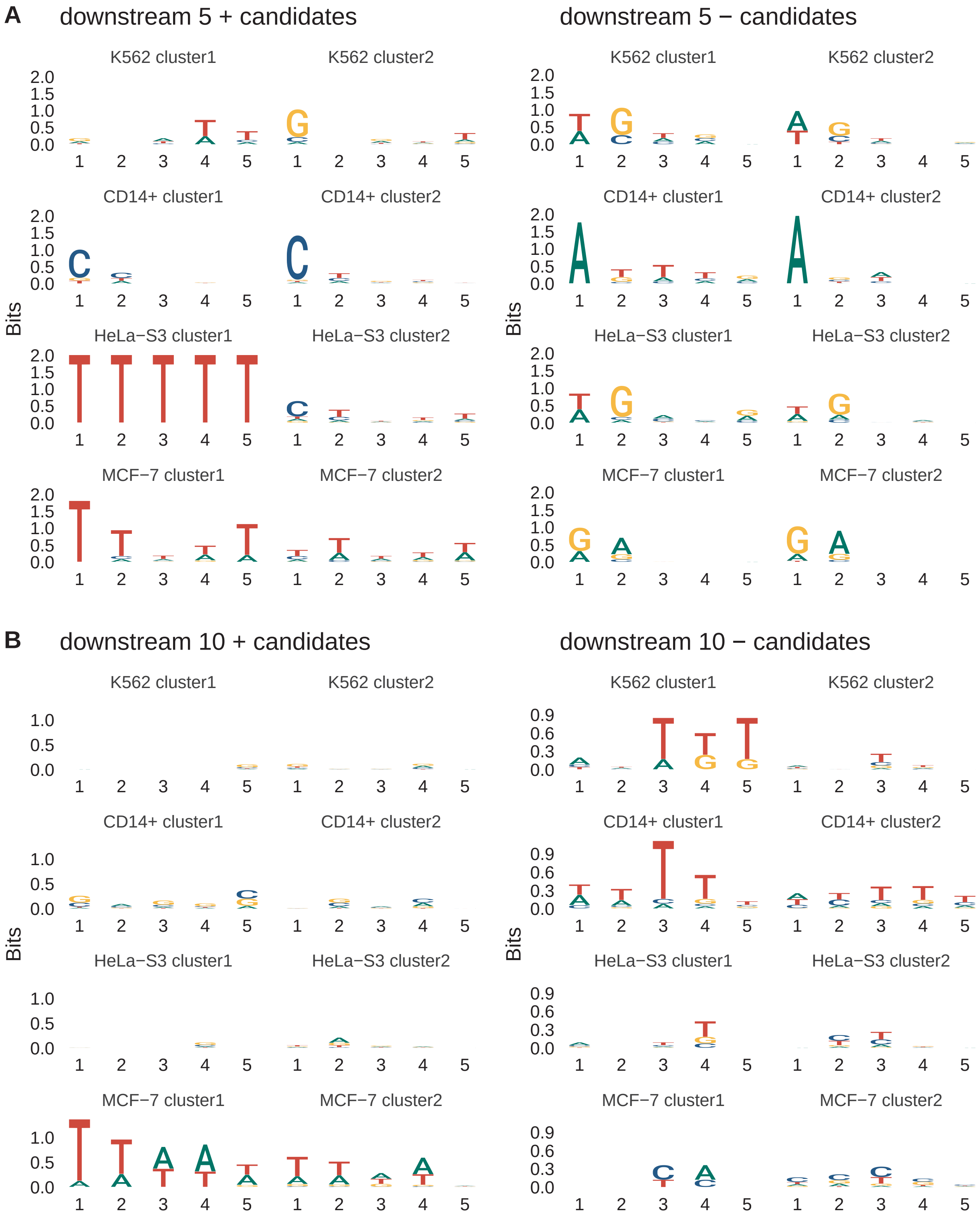


Supplementary Figure S15: **A & B**.Comparison of enrichment logos illustrating the top positive or negative 5-mers in the downstream 5-nucleotide and 10-nucleotide regions along gene bodies for four cell lines.

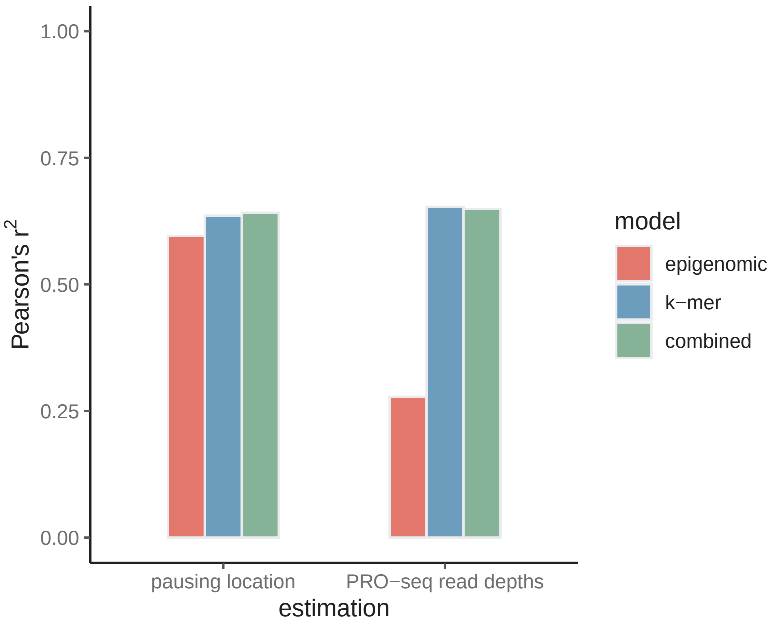


Supplementary Figure S16: The predictive performance of various model versions, such as epigenomic, $k$-mer, and combined epigenomic and $k$-mer models, is assessed using held-out data. Prediction accuracy is measured through the estimation of pausing locations and PRO-seq read counts in 1 kbp regions.
